# Hi-TrAC reveals fractal nesting of super-enhancers

**DOI:** 10.1101/2022.07.13.499926

**Authors:** Yaqiang Cao, Shuai Liu, Kairong Cui, Qingsong Tang, Keji Zhao

**Affiliations:** Laboratory of Epigenome Biology, Systems Biology Center, Division of Intramural Research, National Heart, Lung and Blood Institute, National Institutes of Health, Bethesda, Maryland, USA

**Keywords:** Hi-TrAC, segregation score, active sub-TADs, super-enhancers

## Abstract

Eukaryotic genome spatial folding plays a key role in genome function. Decoding the principles and dynamics of 3D genome organization depends on improving technologies to achieve higher resolution. Chromatin domains have been suggested as regulatory micro-environments, whose identification is crucial to understand the genome architecture. We report here that our recently developed method, Hi-TrAC, which specializes in detecting chromatin loops among genomic accessible regulatory regions, allows us to examine active domains with limited sequencing depths at a high resolution. Hi-TrAC can detect active sub-TADs with a median size of 100kb, most of which harbor one or two cell specifically expressed genes and regulatory elements such as super-enhancers organized into nested interaction domains. These active sub-TADs are characterized by highly enriched signals of histone mark H3K4me1 and chromatin-binding proteins, including Cohesin complex. We show that knocking down core subunit of the Cohesin complex using shRNAs in human cells or decreasing the H3K4me1 modification by deleting the H3K4 methyltransferase *Mll4* gene in mouse Th17 cells disrupted the sub-TADs structure. In summary, Hi-TrAC serves as a compatible and highly responsive approach to studying dynamic changes of active sub-TADs, allowing us more explicit insights into delicate genome structures and functions.

**Highlights:**

- Hi-TrAC detects active sub-TADs with a median size of 100 kb.
- Hi-TrAC reveals a block-to-block interaction pattern between super-enhancers, and fractal structures within super-enhancers.
- Active sub-TADs are disrupted by the knockdown of RAD21.
- Active sub-TADs interaction densities are decreased by the knockout of *Mll4*.

## INTRODUCTION

Proper multi-scale chromatin spatial folding is essential for eukaryotic genome packaging and carrying out biological processes such as DNA replication (1,2), immunoglobulin heavy chain V(D)J recombination (3-5), and cell differentiation (6-9). Three-dimensional genome structural disorganization is associated with the pathogenesis of cancers and developmental diseases (10-13). Understanding of the multilayer 3D genome is evolving with the development of powerful new technologies, including super-resolution microscopy imaging methods and chromosome conformation capture (3C)-derived methods based on proximity ligation and high-throughput sequencing (14-16). The initial version of genome-wide 3C method, Hi-C, led to the discovery of large compartments at the megabase scale with a low resolution of 100 kb (17). With deeper sequencing, topologically associating domains (TADs) at the sub-megabase scale were identified at a better resolution of 40 kb (18-20). With the improved in situ Hi-C method and with much deeper sequencing, chromatin loops and hierarchically nested sub-TADs were identified at much higher 1-5 kb resolution (21). Recently, Hi-C heir Micro-C was able to achieve a resolution of the single nucleosome level, using billions of sequencing reads (22,23). Accompanied by the evolution of Hi-C and its variants, other sequencing-based methods with varying principles, such as ChIA-PET (24), GAM (25), SPRITE (26), and Trac-looping (27), have all been able to validate the existence of high-order structures of compartments, domains, and loops, advancing our understandings of dynamics of the 3D genome.

Since the initial identification of TADs, domain-centric analysis has remained essential in interpreting genome-wide sequencing data and elucidating 3D genome organization and regulation. Features of TADs, including their existence in a range of species (18,19,28,29), boundaries associated with CTCF and Cohesin (18,30-32), formation mechanism through loop-extrusion model (33), and regulatory role in gene expression (11,13) have been extensively documented with Hi-C data (34,35). Therefore, with a physical size of around several hundred nanometers (16,36), TADs are considered as the architectural chromatin units that define regulatory landscapes, framing the microenvironment for enhancer-promoter interactions. Our recent work, Hi-TrAC (37), provides high-resolution maps of interactions between accessibility sites as loops and generic chromatin interactions analogous to Hi-C and variants as domains (38). The loop-centric analysis was performed comprehensively (37), while domain-centric analysis was not included.

By explicitly examining the domain features revealed by modestly-sequenced Hi-TrAC data, we found that it can detect domains around 100 kb in size. These domains show cell specificity, harbor cell identity genes and super-enhancers; they contain enriched ChIP-seq signals of active histone modification marks and transcription factors including H3K4me1 and SMC3. We observed self-similar domain structures within the super-enhancers and their component enhancers. We found that super-enhancers interact with each other with a unique block-to-block pattern when they are in the same domain. Knocking down RAD21 or deleting the H3K4 methyltransferase *Mll4* gene decreased interactions in these domains. Collectively, our results demonstrate that Hi-TrAC is a highly robust and compatible method for studying active domains at a high resolution, even with limited sequencing depth.

## RESULTS

### Segregation scores reveal TADs-like domains from Hi-TrAC data

Insulation scores (28,55) are widely used to obtain domain boundaries from Hi-C and its variants data by transforming two-dimensional contact matrix into one-dimensional signals. Based on the insulation scores, the resulting domains are dependent on the critical parameters of resolution, sliding window size, and the cutoff of insulation score to define the boundaries and combinations of boundaries to dictate domains. Especially when comparing different samples, parameter tuning may be further needed. In addition to these constraints, we found that the insulation scores calculated in Hi-TrAC cannot accurately indicate the boundaries with the lowest local values. To overcome such limitations, we proposed the segregation score to identify putative domains in Hi-TrAC data (**Methods**) (**Figure S1A**). These two approaches share the same idea of transforming two-dimensional matrix into one-dimensional signals. Unlike the insulation score, with which the local lowest score indicates the domain boundary, positive segregation score regions are stitched together as the domain region for Hi-TrAC data (**Figure S1A-B**). No extra cutoff for the segregation score is needed. Two other metrics were required to define a putative Hi-TrAC domain: (1) more interacting paired-end tags (PETs) within the putative domain than the PETs with only one end located, which is the natural definition of the interaction domain, termed as domain enrichment score (ES) for all following analysis; (2) two-folds more interaction density than chromosome-wide interaction density. To test this method, we performed the domain-calling algorithm based on the segregation score at a resolution of 10 kb in GM12878 Hi-TrAC data. The resulting domains are TAD-like and visually consistent with domains from Hi-C data (21) (**Figure S1B and C**), suggesting that the segregation score is reliable for detecting domains from Hi-TrAC data. We also noticed more explicit inner structures than Hi-C data in these Hi-TrAC domains (**Figure S1B and C**), demonstrating the ability of Hi-TrAC data to reveal fine-scale structures of smaller domains at higher resolution.

### Hi-TrAC detects active sub-TADs

To see whether modestly-sequenced Hi-TrAC data (GM12878: 116 million unique intra-chromosomal PETs; K562: 95 million unique intra-chromosomal PETs) can reveal smaller domains at an even better resolution than Hi-C, we performed the domain-calling at a resolution of 1 kb. Among the top-ranked Hi-TrAC domains in GM12878 sorted by segregation scores (descend ranking), we found multiple intriguing features (**Figure 1 A and B**): (1) domain sizes are smaller than 200 kb; (2) top domains contain super-enhancers; (3) a block-to-block interaction pattern exists within and between super-enhancers (in contrast to the typical dot-to-dot loop Hi-TrAC pattern) (37); and (4) top-ranking domains harbor important functional genes in B cells. Since the GM12878 cell line is a B lymphocyte cell transformed by the Epstein-Barr virus, these results suggest that domains may harbor the cell identity genes for GM12878, which were exemplified by the genomic locus of B cell proliferation- and differentiation-related genes such as *BTG2* (56) and *IKZF3* (57) (**Figure 1 A and B**). It was noticed that strong boundaries preferentially insulate super-enhancers in Hi-C data (58), and super-enhancers are highly enriched among TAD triples in GAM data (25). Thus, the block-to-block interaction patterns observed in our Hi-TrAC data provide compelling evidence in support of individual super-enhancers as domain boundaries.

**Figure 1.**
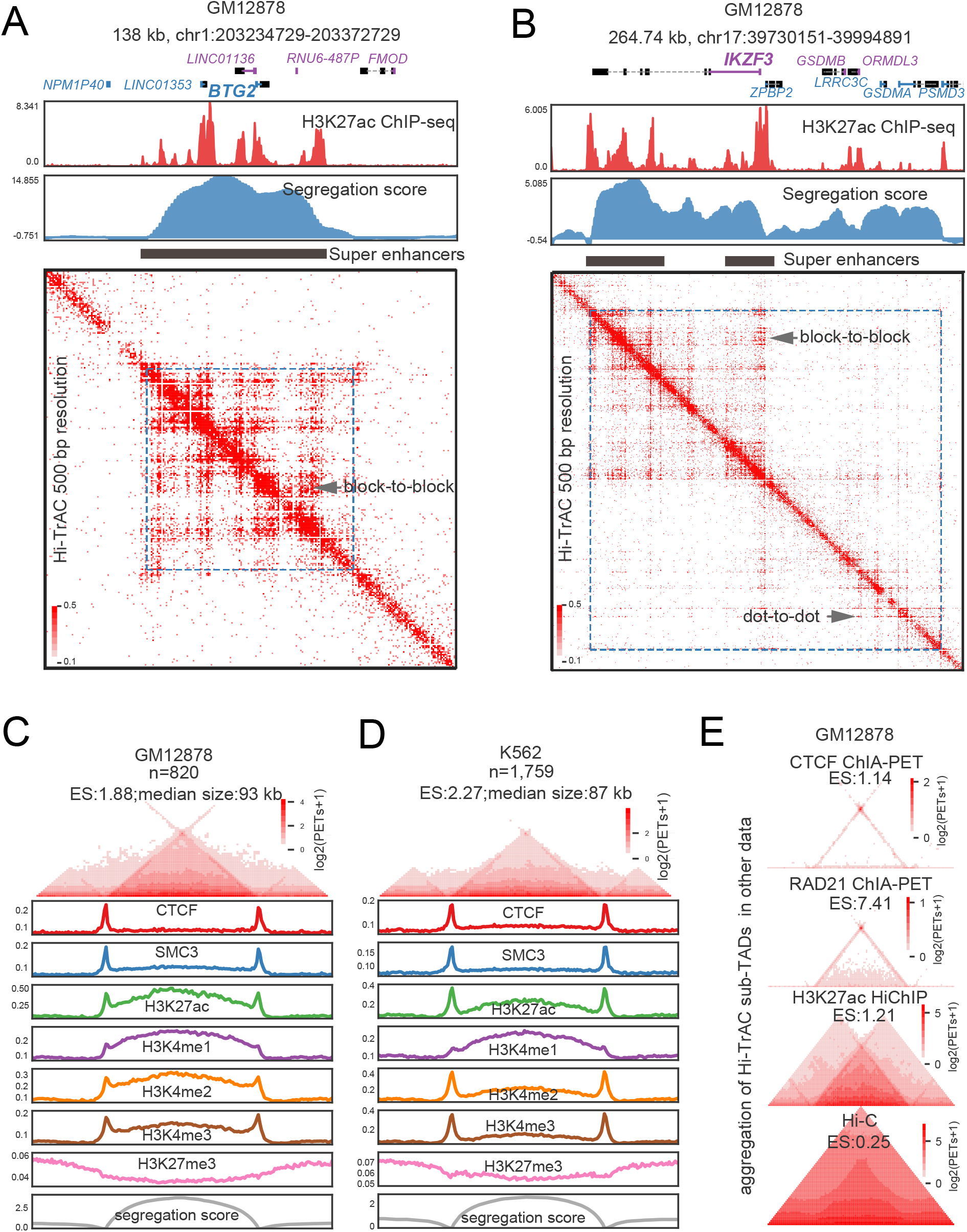
Identification of active sub-TADs by Hi-TrAC. (A) Example of active sub-TAD containing the *BTG2* gene locus detected by Hi-TrAC in GM12878 cells. The cLoops2 callDomains module called domains at 1kb resolution (**Methods**). The first exon of a gene and its name in the positive strand are indicated by blue color, and the first exon of a gene and its name in the negative strand are indicate by purple color. ENCODE (70) H3K27ac ChIP-seq profiles and the Hi-TrAC domain segregation scores are displayed below the genomic annotations. The filled black rectangle indicates the super-enhancers within the active sub-TADs. Hi-TrAC interaction matrix was shown at 500 bp resolution. The interaction domain was marked as blue dotted frames on the heatmap. The plot was generated by the cLoops2 plot module. (B) Example of active sub-TAD containing the *IKZF3* gene locus detected by Hi-TrAC in GM12878 cells. (C) Aggregation analysis of 820 active sub-TADs identified from the Hi-TrAC data in GM12878 cells (**Supplemental Table 2**). Together with 0.5-fold neighboring upstream and downstream regions, all the Hi-TrAC active sub-TADs were aggregated for visualization (**Methods**). ES stands for enrichment scores of interacting PETs within the domains compared to the PETs with one end in the domain and the other outside the domain. The enrichments of various ChIP-seq signals were calculated based on ENCODE ChIP-seq data (70). The analysis was performed with the cLoops2 agg module. (D) Aggregation analysis of 1,759 active sub-TADs identified from the K562 Hi-TrAC data (**Supplemental Table 2**). (E) Aggregation analysis of Hi-TrAC active sub-TADs with CTCF ChIA-PET data (24), RAD21 ChIA-PET data (66), H3K27ac HiChIP data (67) and in situ Hi-C data (21) in GM12878 cells.

In total, we identified 820 domains with a median size of 93 kb in GM12878 (**Figure 1C** and **Supplemental Table 2**) and 1,759 domains with a median size of 87 kb in K562 (**Figure 1D** and **Supplemental Table 2**). As expected, both chromatin domain-boundary associated factors CTCF (18) and Cohesin complex core subunit SMC3 (59,60) have enriched bindings at Hi-TrAC domain boundaries (**Figure 1C** and **D**), indicating their roles in domain establishment or maintenance. As these domains were identified in Tn5 accessible chromatin regions, they carried expected active chromatin features such as elevated H3K4me1, H3K4me2, H3K4me3, and H3K27ac signals inside the domain, as well as decreased H3K27me3 signals compared to flanking regions. Interestingly, H3K4me3 signals in both GM12878 and K562 are enriched at the boundaries, consistent with the observation that domain boundaries are also enriched with promoters of actively transcribed genes, suggesting that certain promoters may also have boundary function (22,23,61,62). The median size of sub-TADs is typically around 180 kb (21), and insulated neighborhoods, one of its sub-classes, has a similar size of approximately186 kb (63). Previously identified supercoiling domains are around 100 kb (64), coinciding with the size of recently identified chromatin nanodomains through super-resolution microscopy in single cells (14,65). This suggests that most enhancer-promoter interactions are constrained in domains of a similar size. Even though the domain sizes detected by Hi-TrAC have a much shorter length of around 100 kb, smaller than 180 kb of sub-TADs, and more similar to supercoiling domains and chromatin nanodomains, they still fall within the same size magnitude. Additionally, Hi-TrAC domains were identified in a similar way to sub-TADs via computational analysis of sequencing data. Therefore, we coined them active sub-TADs.

We further validated Hi-TrAC active sub-TADs using the aggregation analysis of CTCF ChIA-PET data (24), RAD21 ChIA-PET data (66), and H3K27ac HiChIP data (67), but not the in situ Hi-C data (21) due to its limited resolution. The aggregated dot-to-dot interaction patterns at the active sub-TADs boundaries observed from the CTCF and RAD21 ChIA-PET data indicate that Hi-TrAC active sub-TADs may also be formed through the loop extrusion model (33,68), which is similar to the establishment of insulated neighborhoods (63,69), albeit at a smaller scale. We compared these data around a Hi-TrAC active sub-TAD covering the genomic locus of the *TMSB4X* gene in GM12878 (**Figure S2A** and **B**), further validating the observation that other techniques may detect the active sub-TADs but not as straightforward as Hi-TrAC, both for the boundaries and inside structures. Together, our results indicate that Hi-TrAC is more able to capture small active chromatin domains than other methods.

### Hi-TrAC reveals fractal nesting structures within super-enhancers

To follow up on our observation that the active sub-TADs with the high segregation scores in GM12878 contain super-enhancers (**Figure 1A** and **B**), we performed more analysis for the association between Hi-TrAC active sub-TADs and super-enhancers. We found that around 80% (204 of a total 257) and 56% (413 of a total 731), respectively, super-enhancers in GM12878 and K562 cells overlapped with active sub-TADs (**Figure 2A**). Additionally, active sub-TADs with super-enhancers have higher segregation scores, enrichment scores (defined as **Figure S1A**), and interaction densities than the active sub-TADs without super-enhancers (**Figure 2B**), indicating these domains are more insulated and interactive. Even within the same active sub-TAD, there are block-to-block interaction patterns between composition enhancers from both the same and different super-enhancers (**Figure 2C**). Zooming in on the domains at a 500 bp or 200 bp resolution revealed a fractal nesting structure within the individual composition enhancers of each super-enhancer (**Figure 2D** and **E**). This inner nesting structure has not been directly observed before through contact matrix heatmaps due to limitations of resolution (**Figure S3**). Binding of SMC3 was enriched at some boundaries of the inner nesting structure and thus Cohesin may be important for the genomic folding within super-enhancer composition enhancers (**Figure 2D**).

**Figure 2.**
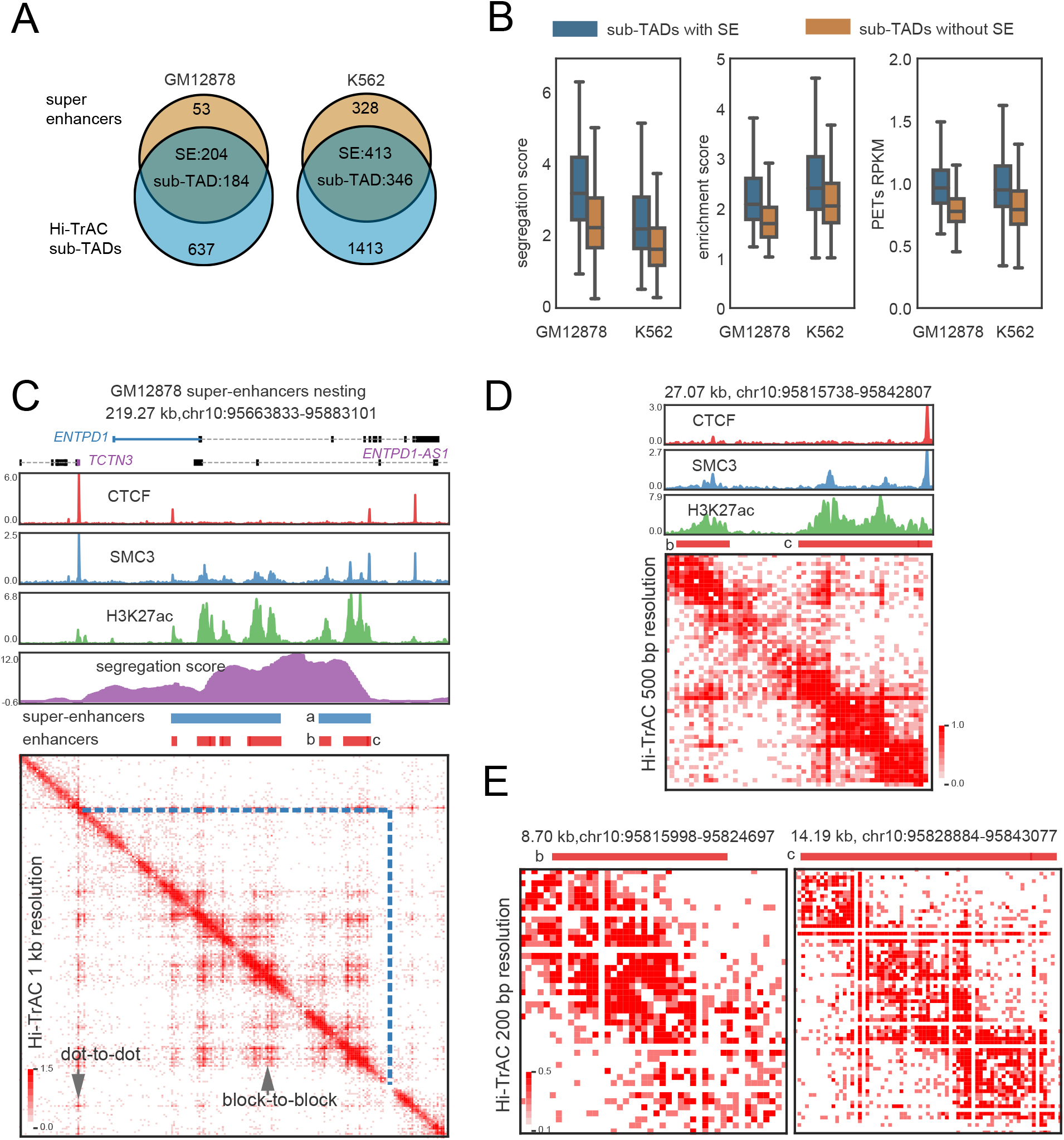
Hi-TrAC reveals the inner structure of super-enhancers. (A) Overlaps of active sub-TADs and super-enhancers in GM12878 and K562 cells. (B) Comparisons of segregation score, enrichment score, and interaction densities between active sub-TADs with and without super-enhancers. (C) Example of active sub-TAD containing the *ENTPD1* gene locus detected by Hi-TrAC in GM12878 cells. Super-enhancer marked with “*a”* and its composition enhancers marked with “*b”* and “*c”* are further visualized. (D) Zoom-in visualization of the inner structure of super-enhancer marked with “*a”* in panel (C). (E) Zoom-in visualization of the inner structure of enhancers marked with “*b”* and “*c”* in panel (C).

### Active sub-TADs associated epigenetic features

Over 60% of active sub-TADs contained fewer than two genes (**Figure 3A**), which exhibited significantly higher expression levels than other genes (**Figure 3B**). This implies that a substantial number of highly transcribed genes are regulated within their own gene-specific domains. Gene ontology (GO) analysis showed that the genes harbored in active sub-TADs were related to regulation of T cell activation, regulation of hemopoiesis, lymphocyte differentiation, and positive regulation of cytokine production, indicating that these domains may mark specific GM12878 cells’ cellular identity and functions (**Figure 3C**). Together, these results suggest that active sub-TADs may play important roles in regulating and determining cell function.

**Figure 3.**
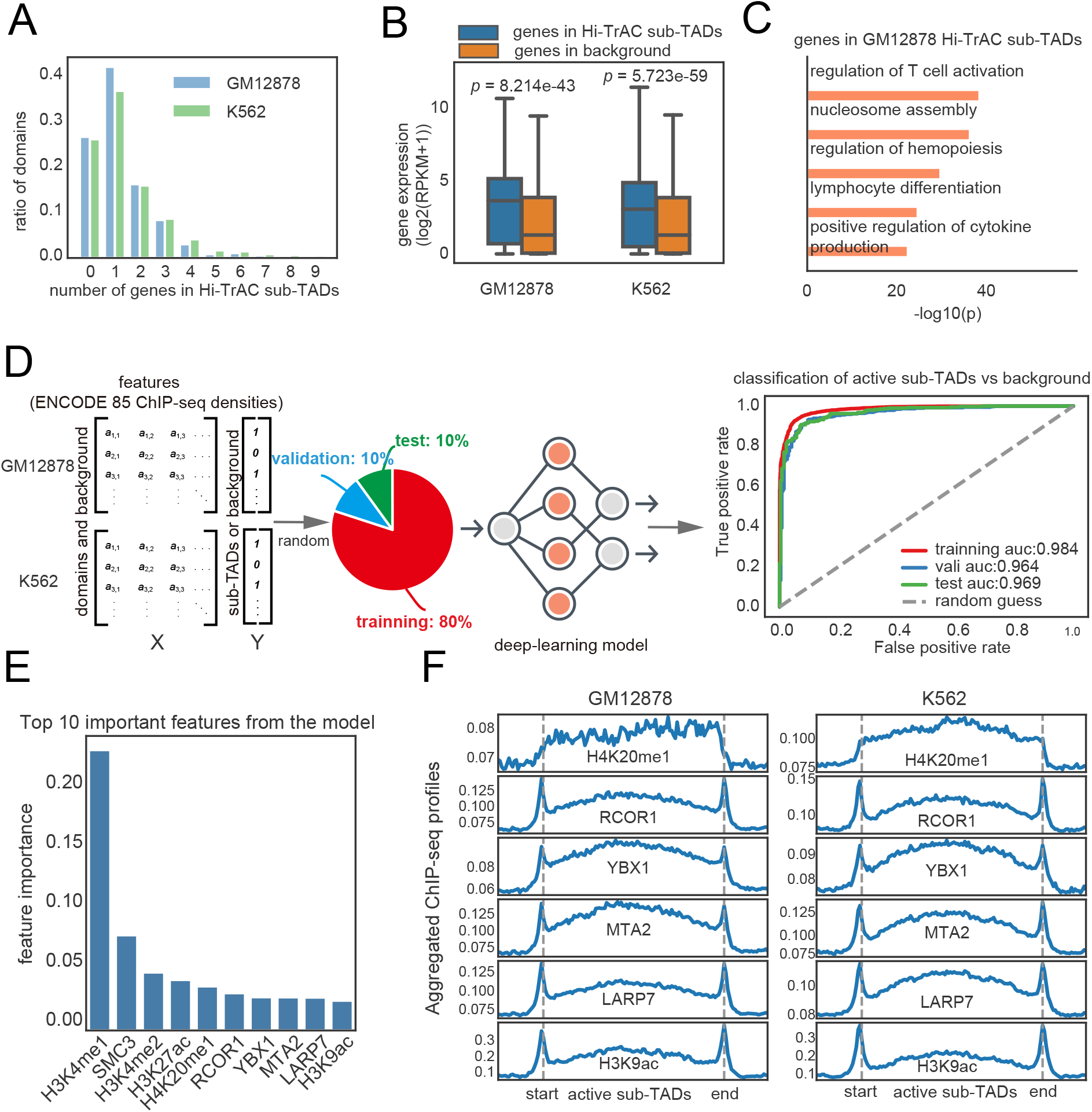
Properties of Hi-TrAC detected active sub-TADs. (A) Distribution of numbers of genes in an active sub-TAD. (B) Distribution of expression levels for the genes located in the active sub-TADs and background regions. RNA-seq data were obtained from GSE30567 (88). Background regions were defined as the same sized regions as sub-TADs but located either upstream or downstream, not overlapped with any active sub-TADs. Wilcoxon rank-sum test was used to calculate *P*-values. (C) Gene ontology (GO) analysis for genes located within the Hi-TrAC active sub-TADs in GM12878 cells. The top 5 enriched GO terms were shown. (D) Scheme and receiver operating characteristic (ROC) curves for the performance of classification model for the identification of important features of active sub-TADs using ENCODE ChIP-seq data for GM12878 and K562 (**Methods**). All data were separated into training, validation, and test sets (8:1:1), with training data used to train the model, validation data used to select the model, and test data used to evaluate the model’s final performance. (E) Top ten most important features associated with the active sub-TADs. (F) Aggregation analysis for ChIP-seq signals of the top important features in the active sub-TADs and nearby background regions.

To further study the epigenetic features associated with active sub-TADs, which may provide information on the potential regulators for these active sub-TADs, we quantified the enriched signals of ENCODE (70) ChIP-seq data of 85 factors (shared between GM12878 and K562) in active sub-TADs and flanking background regions (**Figure S4A**) in both GM12878 and K562 cells (**Supplemental Table 1**). These 85 factors included histone modifications and transcription factors (TFs). We implemented a deep learning model to investigate whether epigenetic markers and TFs ChIP-seq signals have the classification power to distinguish the active sub-TADs from the flanking backgrounds (**Figure S4B**) (**Methods**). If the model is accurate, it should be able to capture the important latent features and assign the high-value weights for them, which could then reveal important regulatory factors. After training, the model was able to classify active sub-TADs against background based on 1D ChIP-seq information (**Figure S4C and Figure 3D)**. Even for the test data (10% of all data) that had never been previously used in model training or validation, the classification accuracy remained reasonably high (0.911: the percentage of correctly classified items in the total item count) (**Figure S4C**). The areas under the receiver operating characteristic curve (AUCs) were higher than 0.96 throughout training, validation, and test datasets, indicating that the model was properly trained, not overfitted, and highly reliable (**Figure 3D**). Equipped with high accuracy, the model can reliably reveal important features associated with active sub-TADs (**Figure S4D**) (**Methods**). According to the classification model, H3K4me1 is the most important feature in classifying active sub-TADs against background (**Figure 3E**). (44,71)SMC3 ranks top 2 as a known chromatin domain associated factor. Other top features including active histone marks H4K20me1(top 5), H3K9ac (top 10) and transcription factors RCOR1, YBX1, MTA2, and LARP7 exhibited elevated signals in the active sub-TADs and at the boundaries (**Figure 3F**). (45,72)However, it is unclear whether the enrichment of these features in these active sub-TADs is causative or simply correlative (**Figure 3F**).

### Cell-specific active sub-TADs harbor cell identity genes

To study the cell specificity of active sub-TADs, we compared active sub-TADs detected in GM12878 and K562 cells. We identified 538 (65% of a total 820) GM12878 specific active sub-TADs, and 818 (46% of a total 1,759) K562 specific active sub-TADs based on the difference in segregation scores (**Figure 4A, Supplemental Table 3, Methods**). Cell-specific active sub-TADs in their respective cells showed higher signals of the top ten important features identified from the classification model (**Figure 4A**). The identification of these cell-specific sub-TADs was validated using differential aggregation analysis of the unbiased Hi-C data, and feature-selective H3K27ac HiChIP and RAD21 ChIA-PET data (**Figure 4B**). Our results also show that genes located in cell-specific active sub-TADs have higher expression levels in their corresponding cell type (**Figure 4C)**. Consistent with a B cell transformed cell line, the top enriched KEGG terms for the genes in the cell-specific active sub-TADs in GM12878 cells contained genes involved in the B cell receptor signaling pathway (log(*P*-value) =-18.2, HOMER). For K562, a chronic myelogenous leukemia cell line, one of the most enriched GO terms was myeloid leukocyte mediated immunity (log(*P*-value) =-17.7, HOMER). This was further exemplified by *PLCG2*, a gene critical for B cell signaling (73), which had a super-enhancer and was specifically expressed in an active sub-TAD only detectable in GM12878 cells. The leukemia-associated gene *PIM1* (74), expressed only in K562, was localized to a cell-specific active sub-TAD containing a super-enhancer spanning almost the entire sub-TAD (**Figure 4D**). Although the data from other methods, such as Hi-C and H3K27ac HiChIP, could validate the cell specificity of sub-TADs using aggregate analysis (**Figure 4B**), they were unable to achieve high resolution in interaction heatmaps of specific genomic loci as shown by Hi-TrAC (**Figure S5**). In summary, our results demonstrate that the explicit cell-specific active sub-TADs detected by Hi-TrAC are chromatin structures critical to regulating cell identity and activity.

**Figure 4.**
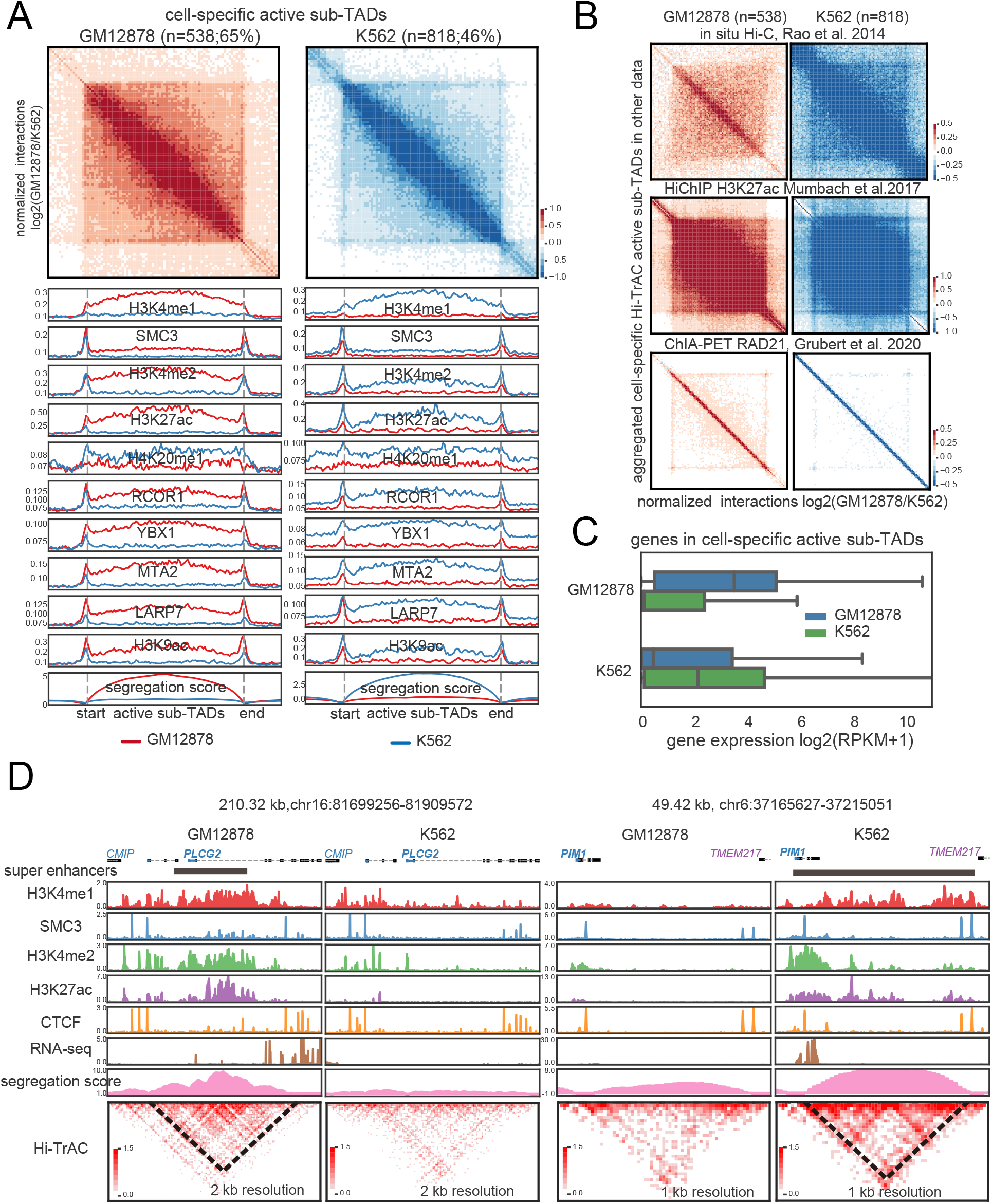
Hi-TrAC detects cell-specific active sub-TADs. (A) Aggregation analysis of cell-specific active sub-TADs in GM12878 and K562 cells (**Supplemental Table 3**). The top ten most important features identified from the classification model were also shown. (B) Aggregated domain analysis for Hi-TrAC detected cell-specific active sub-TADs using in situ Hi-C (top panel), H3K27ac HiChIP (middle panel), and RAD21 ChIA-PET data (lower panel). (C) Distribution of expression levels for the genes located within the cell-specific sub-TADs identified by Hi-TrAC. The blue box shows the genes of GM12878 cell-specific sub-TADs, and the green box shows that of K562, respectively. (D) Examples of GM12878 and K562 specific active sub-TADs.

### Active sub-TADs are disrupted by knocking down RAD21

To test whether CTCF or Cohesin contributes to the maintenance of active sub-TADs, we further analyzed the Hi-TrAC and Hi-C data from K562 cells knocked down of CTCF or RAD21, a major cohesin subunit, using shRNAs. The results from the Hi-TrAC and Hi-C samples showed that knocking down RAD21 substantially impaired chromatin interactions up to hundreds of kb (**Figure 5A** and **B**), suggesting that the changes occur mainly at domain levels, including TADs and sub-TADs. Additionally, the changes in interaction distance were consistent with previous results from Cohesin-depleted Hi-C data in which loop domains were eliminated but compartment domains remained (75). However, knocking down CTCF did not show the impairment of loop domains resulting from knocking down RAD21 (**Figure 5A** and **B**). Taken together, the changes in aggregated segregation scores from Hi-TrAC data (**Figure 5C**) and aggregated differential interaction contact matrix from both Hi-TrAC and Hi-C data **(Figure 5D**) indicated that knocking down RAD21 leads to global disruption across Hi-TrAC sub-TADs. Analysis of segregation scores and interaction matrix from Hi-TrAC data revealed that knocking down resulted in decreased intra-domain interactions (**Figure 5C**) and noticeably blurred domain boundaries (**Figure 5D**). Compared to the shRNA control sample, significantly impaired active sub-TADs by knocking down CTCF, RAD21, or both CTCF and RAD21 were called and checked for overlaps with each other (**Figure 5E**) and overlaps with super-enhancers (**Figure 5F**). This shows that RAD21 plays a much more critical role than CTCF in maintaining the active sub-TADs and the potential role of super-enhancers in chromatin structure. At the leukemia oncogene *LMO2* (76) gene locus (**Figure 5G**), we found that Hi-TrAC is sensitive enough to detect changes in chromatin interactions for active sub-TADs in the same cell type under perturbation. Additionally, the resulting five-fold decrease in LMO2 expression suggests that sub-TADs may also play an important role in regulating gene expression (control RPKM: 13.5 vs. RAD21 KD RPKM:2.3, averages of two replicates, data included in the supplemental table of (37)). In T cell acute lymphoblastic leukemia, the proto-oncogenes *LMO2* was found to be silent and insulated from active enhancers, and removal of the boundary not only enhanced its chromatin interactions with distal region, but also activated its expression in HEK-293T cells (12). In summary, our results collectively demonstrate the critical role of RAD21 in maintaining the sub-TADs detected by Hi-TrAC.

**Figure 5.**
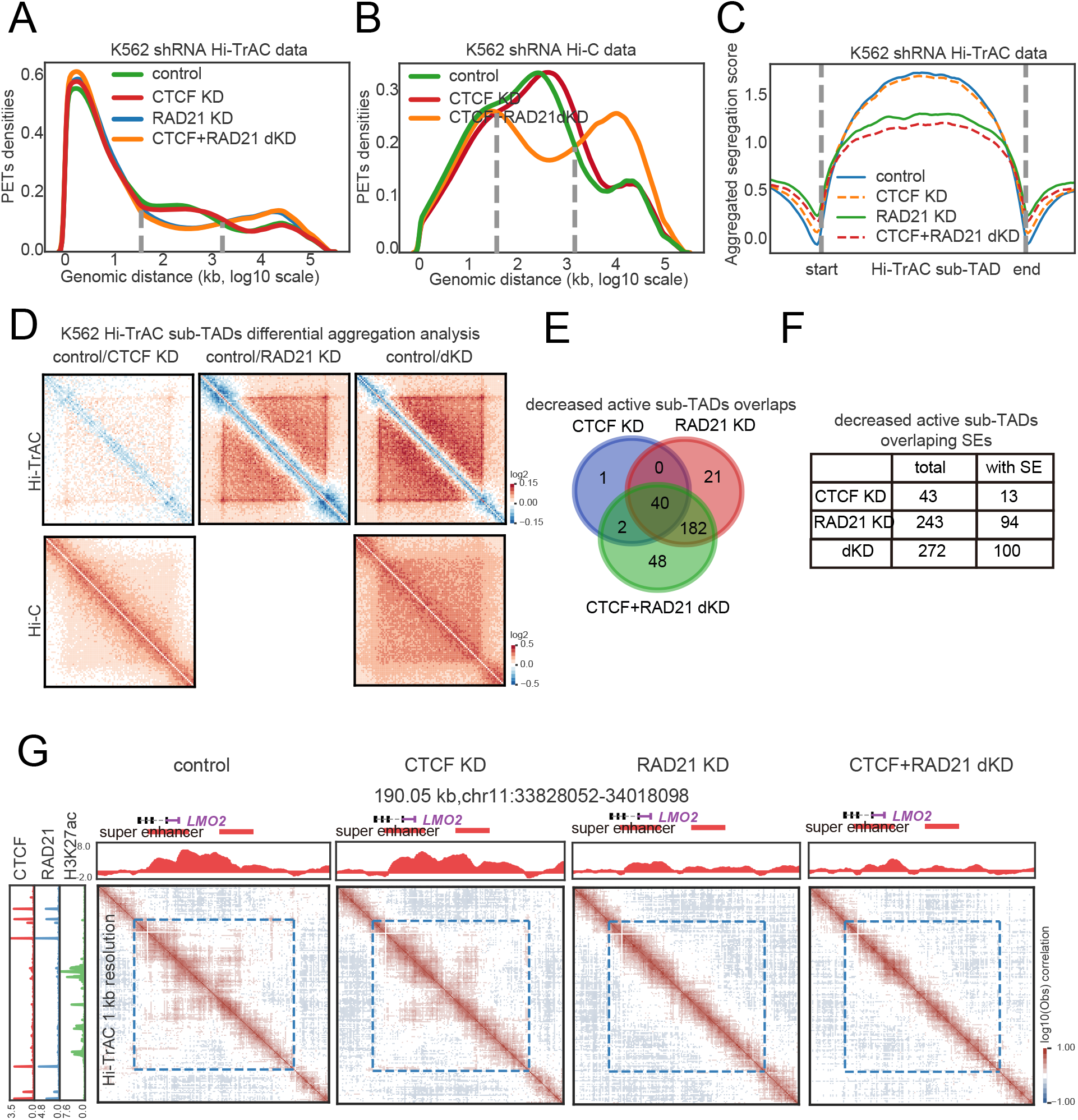
Active sub-TADs are disrupted by knockdown of RAD21. (A) Distribution of interacting PETs frequencies against genomic distances of Hi-TrAC data in wild-type and various knockdown K562 cells. KD: knockdown; dKD: double knockdown. (B) Distribution of interacting PETs frequencies against genomic distances of Hi-C data in wild-type and various knockdown K562 cells. (C) Aggregation analysis of segregations scores of sub-TADs identified from Hi-TrAC data after knocking down CTCF or RAD21 either alone or in combination in K562 cells. (D) Aggregation analysis of interaction matrix of sub-TADs identified from Hi-TrAC data after knocking down CTCF or RAD21 either alone or in combination in K562 cells with Hi-TrAC (upper panel) or Hi-C data (lower panel). (E) Overlaps of significantly changed K562 Hi-TrAC active sub-TADs after knocking down CTCF or RAD21 alone or in combination. (F) Overlaps of significant changed K562 Hi-TrAC active sub-TADs with super-enhancers. (G) Example of disrupted active sub-TADs in K562 by knocking down RAD21 at the *LMO2* gene locus.

### Interactions of active sub-TADs are decreased by deleting *Mll4* in mouse Th17 cells

The prediction model suggested that H3K4me1 is the most enriched feature associated with the Hi-TrAC active sub-TADs (**Figure 3E**), and thus we decided to further investigate whether H3K4me1 contributes these structures. MLL4, also known as KMT2D, is a primary H4K4 mono- and di-methyltransferase in mammalian cells and its deletion significantly decreases H3K4me1 levels on enhancers in T cells (44). Thus, we performed RNA-seq, ChIP-seq for H3K4me1, H3K4me3 and H3K27ac, and Hi-TrAC analyses of T helper 17 (Th17) cells from wild-type and *Mll4* conditional knockout mice (**Supplemental Table 5**). The specific knockout of *Mll4*’s exon was validated by the RNA-seq data (**Figure S6A**). While the global patterns of the histone marks were similar between wild-type and *Mll4* deletion cells (**Figure S6B)**, enhancers showed significantly decreased histone modification signals in the *Mll4* deleted cells (**Figure S6E, Supplemental Table 6**). Based on the Hi-TrAC interaction signals, wild-type and *Mll4* deleted cells were clustered into different groups with either 1Kb or 5Kb resolution (**Figure S6C**). With limited sequencing depth, both pooled wild-type and *Mll4* KO Hi-TrAC data show the highest estimated genome-wide resolution as 1 kb (**Figure S6D**). By examining different regulatory regions, we found that chromatin interactions showed the highest decreases at putative enhancers (**Figure S6F**). By randomly checking the genomic regions, we noticed H3K4me1 and H3K4me3 marks aligned well at TAD-like regions, and decreased interaction densities may happen at around 200 kb scale (**Figure S6G**). The size coincided with the size of active sub-TADs, indicating that Hi-TrAC has the detection sensitivity for individual active sub-TAD with subtle interaction changes.

We called active sub-TADs from Th17 cells based on the wild-type Hi-TrAC data (**Methods** and **Supplemental Table 7**). The *Rorc* locus, encoding the Th17 master transcription factor RAR-related orphan receptor gamma (RORγt) (78), was located in a ∼ 100 kb active sub-TAD (**Figure 6A**), which is consistent with our observation that active sub-TADs harbor cell identity genes in human cells. In total, we identified 1,427 active sub-TADs with a median size of 82 kb (**Figure 6B**). KEGG enrichment analysis for the genes contained within these active sub-TADs revealed interesting terms including *Systemic lupus erythematosus* (SLE), *Th17 cell differentiation*, and *T cell receptor signaling pathway*. These terms are related to Th17 cell differentiation, function, or potential diseases (79-81), further supporting our observation in human cells that active sub-TADs may play a regulatory role in cell functions.

**Figure 6.**
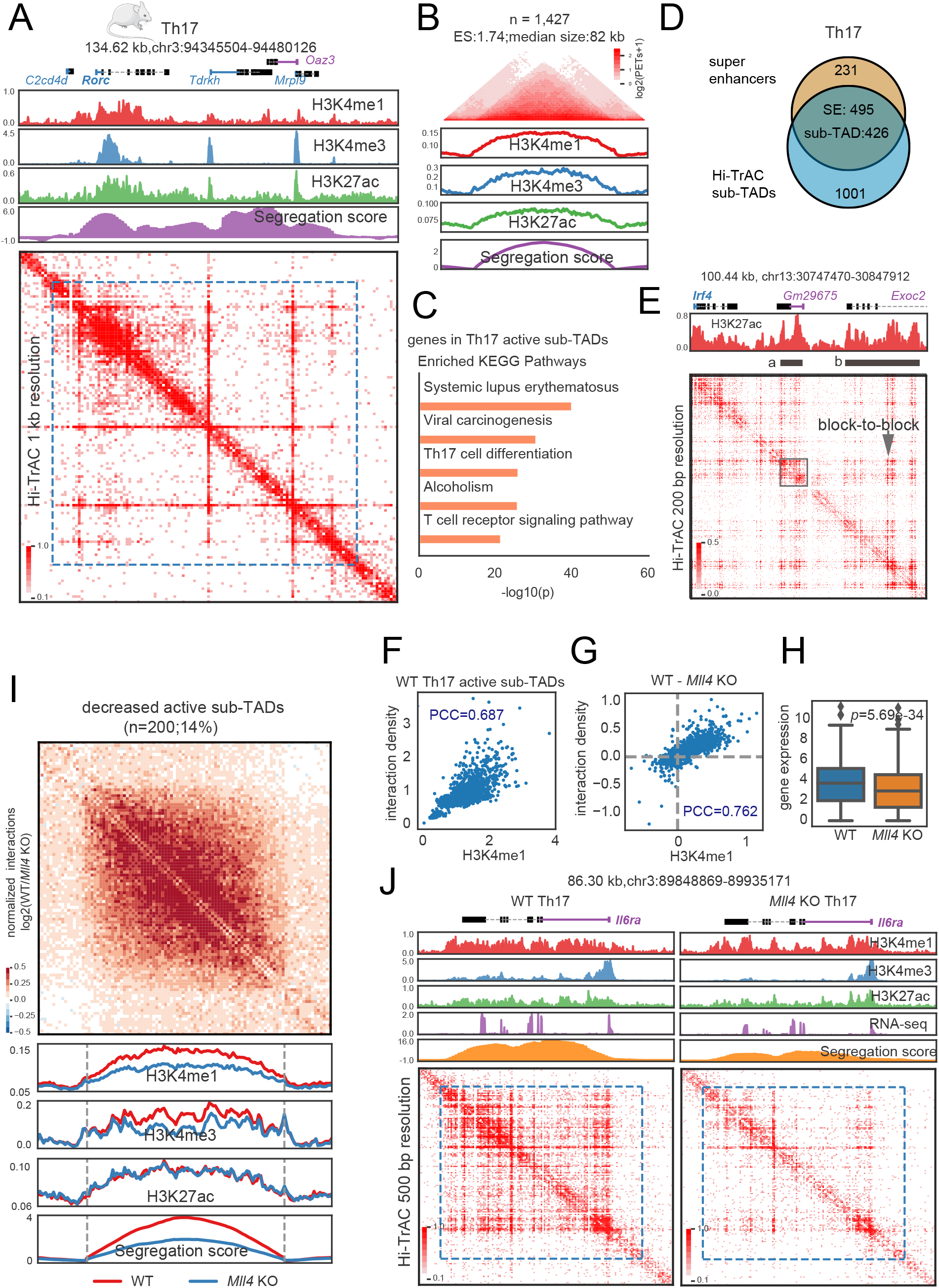
Interactions within active sub-TADs are decreased by deletion of *Mll4* in mouse Th17 cells. (A) The *Rorc* genomic locus, encoding a master transcription factor in Th17 cells, is located in an active sub-TADs. ChIP-seq and Hi-TrAC data were generated in this study (**Supplemental Table 5**). ChIP-seq signals were the average values from two replicates, and Hi-TrAC data were pooled from three biological replicates (**Supplemental Table 5**). (B) Aggregation analysis of 1,427 active sub-TADs identified from Hi-TrAC data in mouse Th17 cells (**Supplemental Table 7**). (C) KEGG terms enrichment analysis for genes located within the Hi-TrAC active sub-TADs in mouse Th17 cells. Only the top 5 enriched terms were shown. (D) Overlaps of active sub-TADs and super-enhancers in mouse Th17 cells. (E) Example of active sub-TAD containing the *Irf4* gene locus detected by Hi-TrAC showing block-to-block interaction pattern between super-enhancers. Two super-enhancers were marked as “a” and “b”. Super-enhancer “a” was highlighted with a gray box to show its inner structure. (F) Correlation analysis between H3K4me1 ChIP-seq signal intensities and Hi-TrAC interaction densities for wild-type Th17 cells active sub-TADs. PCC stands for Pearson Correlation Coefficient. (G) Correlation analysis between the changes in H3K4me1 ChIP-seq signal intensity and Hi-TrAC interaction density in active sub-TADs after deletion of *Mll4* Th17 cells. (H) Distribution of expression levels for the genes located within the active sub-TADs. (I) Aggregation analysis of significantly decreased active sub-TADs based on segregation scores in Th17 cells comparing wild-type and *Mll4* knockout mice (**Supplemental Table 7**). (J) Example of a significantly decreased active sub-TAD harboring the *Il6ra* gene important for Th17 function.

We further called super-enhancers based on H3K27ac ChIP-seq data (**Methods** and **Supplemental Table 6**) and found 495 (68.18%) Th17 super-enhancers overlapped with active sub-TADs (**Figure 6D**), consistent with the result that more than half super-enhancers in human GM12878 and K562 cells were overlapped with active sub-TADs. We also observed block-to-block interactions between the two super-enhancers of the *Irf4* gene locus (**Figure 6E**) and between the inner structures within an individual super-enhancer (**Figure 6E** gray box). The interactions measured by Hi-TrAC in active sub-TADs showed a high correlation with the level of H3K4me1 as the Pearson Correlation Coefficient (PCC) was 0.687 (**Figure 6F**). The decreases in H3K4me1 levels and interactions in the active sub-TADs resulting from *Mll4* deletion were also highly correlated (PCC=0.762, **Figure 6G**). The expression levels of genes in active sub-TADs were decreased by *Mll4* deletion (**Figure 6H**). We further obtained 200 significantly decreased active sub-TADs based on changes of segregation scores (**Figure 6I, Methods**). The H3K4me1 and H3K4me3 levels were decreased in these active sub-TADs but not the H3K27ac (**Figure 6I**). The global pattern was exemplified by the genomic locus of *Il6ra* (**Figure 6J**), whose decreased expression correlates with reduced Th17 response (82) and affects Th17 maintenance (83).

In summary, the Hi-TrAC data from mouse Th17 cells showed consistent results with the data from human cells regarding the size of the active sub-TADs, the high overlap between active sub-TADs with super-enhancers, and enrichment of cell function and identity genes within active sub-TADs. Our results also strongly indicate that H3K4me1 is important for maintaining the interaction densities within active sub-TADs.

## DISCUSSION

Since TADs were first identified, domain-centric analyses of Hi-C data have been widely performed in uncovering the relationship between TADs and the 3D genome folding (34,35). Here we demonstrated that the domain-centric analysis with modest sequenced Hi-TrAC data is a sensitive yet robust method for accurately identifying and studying dynamic changes of active sub-TADs around 100 kb scale with a resolution of 1 kb. The results from our analysis revealed nested domain structures within multiple super-enhancers, which could not be explicitly observed from the interaction matrix heatmaps from current methods such as Hi-C, CTCF ChIA-PET, or H3K27ac HiChIP data. Thus, Hi-TrAC holds a considerable advantage in profiling genome organizations, especially in highly accessible regions at a fine-scale resolution. However, the exact mechanism of why and how the super-enhancers maintain such nested structures remains elusive. It is currently being debated whether TADs reflect probabilistic preferential interactions from bulk cells or stable domains (84-86), and this debate itself could be directly linked to a parallel debate surrounding the existence of super-enhancer inner structures. Future studies of single-cell level data may prove to be a worthwhile approach to address these hypotheses.

Our integration of the ENCODE ChIP-seq data and active sub-TADs using a classification model revealed potentially important factors associated with active sub-TADs. Among these factors, Cohesin and active histone marks of H3K4me1 and H3K27ac were top-ranked, while CTCF was not on the top 10 list. Further, our knockdown experiments supported the role of Cohesin but not CTCF in the maintenance of the active sub-TAD structure, partially validating the model. The model was further validated by deletion of the *Mll4* gene in mouse Th17 cells, which simultaneously decreased the H3K4me1 signals and the interaction density within the active sub-TADs. We also believe that this model will serve as an excellent framework for identifying more factors associated with active sub-TADs and will improve as the ENCODE consortium expands its high-quality datasets (87). However, there are two important caveats in this study to note: 1) It is still the correlation analysis for the signal enrichment at active sub-TADs, which does not indicate any causality. 2) Potential cell-specific factors were excluded as only factors shared between GM12878 and K562 were used to improve the robustness of the model. A potential solution to the first issue is experimental perturbation and validation. For the second issue, a more detailed analysis focused solely on the K562 dataset may prove worthwhile as it has much rich public data, and experimental manipulation of shRNA knockdown or CRISPR/Cas9 knockout is much easier.

With its compatibility and sensitivity in this study, Hi-TrAC has been shown to be an effective method in studying chromatin domains covering regulatory elements at high resolutions, even with modest sequencing depths. We hope that Hi-TrAC will serve as a valuable tool for the 3D genome research community and further our understanding of chromatin domain dynamics in developmental and disease processes.

## METHODS

### Public data and pre-processing

Public data used in this study, including human Hi-TrAC, Hi-C, HiChIP, ChIA-PET, and ChIP-seq, were summarized in our previous work (37). Hg38 alignment BAM files of ENCODE ChIP-seq data were obtained from ENCODE website and summarized in **Supplemental Table 1**. Biological and technical replicates were merged for the same factor, and only unique reads were used for the following analyses. Human gene annotations from GENCODE (39) (gencode.v30.basic.annotation.gtf) were used in any gene-related analysis. Hg38 was used in this study. If human data processed in other genome versions were downloaded, they were always converted to hg38 for analysis. GM12878 and K562 super-enhancers were downloaded from dbSUPER (40) in hg19 and converted to hg38 by UCSC liftOver.

### Segregation score calculation and domain calling from Hi-TrAC data

Domains from Hi-TrAC data were called based on segregation scores. For each bin at the assigned resolution (parameter) from the contact matrix, its upstream and downstream window size (parameter) region is used to construct the contact matrix. The contact matrix is further log_2_ transformed and calculated as a correlation matrix. For the up-right corner sub-matrix of the correlation matrix, all < 0 values are assigned as 0, and the mean value of the matrix is assigned as a segregation score for the bin. After calculating segregation scores for all bins in one chromosome, z-score normalization was performed to the segregation scores, and bins with >0 segregation scores were stitched together as candidate domains. These candidate domains further required an enrichment score (the number of PETs within the domain divided by the number of PETs with only one end within the domain) >=1, and the interaction density is higher than the two folds of chromosome-wide density. The scheme of the algorithm was summarized in **Figure S1A** and implemented as the cLoops2 callDomains module (38). For calling sub-TADs from Hi-TrAC data, bin size was set to 1 kb, and windows size was set to 50 kb, details parameters of -bs 1000 -ws 50000 -strict were used.

### Domain aggregation analysis

For a domain with its 0.5-fold sized neighboring upstream and downstream regions, interacting PETs were grouped into a 100 × 100 bins matrix. An individual enrichment score for a domain is calculated as the number of PETs with both ends located within the domain compared to the number of PETs with only one end located within the domain. The global enrichment score is the mean value of all enrichment scores for individual domains. Heatmap was plotted of the upper triangular matrix of the average matrix by default. The analysis was implemented in the cLoops2 agg module with the option of -domains, and default parameters were used in all related analyses in this manuscript (38).

### Deep-learning model for classification of Hi-TrAC active sub-TADs against background regions with ChIP-seq data

A deep-learning model was implemented based on 85 shared transcription factor or histone modification ChIP-seq datasets between GM12878 and K562 from ENCODE. For the classification of Hi-TrAC active sub-TADs against background regions, background regions were defined as the same sized flanking regions as active sub-TADs but not overlapping with any of them. ChIP-seq reads were quantified as RPKM values for active sub-TADs and background. All data were separated randomly into the training set, validation set, and test set, with a ratio of 0.8:0.1:0.1. The training set was used to train models, and the model was selected by the lowest loss observed for the validation set. The test set was finally used to evaluate the performance of the model. The high accuracy of the classification model should capture the important latent features for distinguishing Hi-TrAC active sub-TADs from the background with ChIP-seq data. Therefore, the estimation for feature importance was performed for each factor by shuffling the RPKM values for all regions 100 times. Then the mean value of decreased accuracy was used as the feature importance. The deep-learning-related functions were powered by the Keras module in TensorFlow (41), and ROC curves were powered by the scikit-learn (42). The scheme of the algorithm was summarized in **Figure S4D**.

### Calling cell- or condition-specific active sub-TADs

Segregation scores of active sub-TADs were quantified both in GM12878 and K562 with the cLoops2 quant module (38), and segregation score difference > 1 was used to get GM12878 specific active sub-TADs. K562-specific active sub-TADs were called in the same way. Significantly lost active sub-TADs after knockdown of CTCF or RAD21, or combined were called in the same way.

### GO terms enrichment analysis

Gene Ontology (GO) terms enrichment analysis for genes was performed by findGO.pl in the HOMER package (43), requiring more than ten overlapping genes in the terms, and there are fewer than 1000 genes in the terms. Only top enriched terms sorted by ascending *P*-values were shown.

### Generation of mouse Th17 cells

*Mll4*^fl/fl^ *CD4*-Cre+ mice were described previously (44). Naive CD4^+^ T cells were purified from the lymph nodes of *Mll4*^+/+^ *CD4*-Cre+ and *Mll4*^fl/fl^ *CD4*-Cre+ mice with EasySep Mouse CD4+ T cell Isolation Kit (StemCell, #19852). Cells were cultured in Th17 differentiation medium (2μg/ml anti-CD28, 10μg/ml anti-IL4, 10μg/ml anti-IFNg, 10μg/ml anti-IL12, 10ng/ml TGFβ, 20ng/ml IL6, 10ng/ml IL1β) on plates coated with anti-CD3 and anti-CD28. Cells were collected by cell sorting with DAPI on 24hrs and 72hrs for RNA-seq, ChIP-seq, and Hi-TrAC.

Hi-TrAC was performed as described previously (37). Briefly, cells were fixed with 1% Formaldehyde at room temperature for 10 minutes. The biotinylated linker for bridging interacting chromatin was inserted into the genome by Tn5. After reverse crosslinking, genomic DNA was purified, and with gaps repaired. After digesting with MluCI and NlaIII restriction enzymes, biotinylated DNA fragments were enriched with streptavidin beads. Universal adapters were ligated to DNA fragments, and the libraries were amplified by PCR with Illumina Multiplexing primers.

ChIP-seq assays were performed as described previously (45,46). In brief, cells were fixed with 1% formaldehyde at room temperature for 10 minutes. Chromatin was sonicated and immunoprecipitated with anti-H3K4me1(ab8895, Abcam), anti-H3K4me3 (17-614, Millipore) and anti-H3K27ac (ab4729, Abcam) antibodies. Purified ChIP DNA was repaired with End-It DNA End-Repair Kit (Epicentre). The library was then indexed and amplified and sequenced on an Illumina platform. For RNA-seq, RNA from 5,000 cells were purified with QIAzol lysis reagent (QIAGEN) and RNeasy mini kit (QIAGEN). The libraries were then constructed following Smart-Seq2 method (47).

### Mouse Th17 ChIP-seq, RNA-seq and Hi-TrAC data analysis

Mouse Th17 ChIP-seq data were mapped to mouse reference genome mm10 with Bowtie2 (v.2.3.5) (48). Only non-redundant reads with MAPQ >=10 were used for the following analysis. Sample-wise genome-wide correlation analysis was performed with multiBigwigSummary bins and parameter -bs 1000 in deepTools2 (v3.3.0) package (49). Peaks were called for each ChIP-seq library first with the cLoops2 callPeaks module (38); specific parameters of -eps 300,500 - minPts 10,20 -sen were used for H3K4me1 datasets, and -eps 150,300 -minPts 10,20 were used for H3K4me3 and H3K27me3 datasets. Overlapped regions from the same histone modification’s biological replicates and wild-type or *Mll4* knockout conditions were compiled together as union sets (**Supplemental Table 6**). The overlapped peaks of H3K4me1, H3K4me3, and H3K27ac were assigned as putative genomic segments with the following criteria (**Supplemental Table 6**): 1) H3K4m3 overlapped H3K27ac peaks at the transcription start site (TSS) as active TSS first; 2) H3K4me3 peaks without H3K27ac peaks at TSS as poised TSS; 3) H3K27ac peaks at TSS distal region as active enhancers; 4) H3K4me1 only peaks at TSS distal region as poised enhancers; 5) finally, other parts of H3K4me1 peaks overlapped with above segments but not totally covered as the flank regions. Super-enhancers were called based on wild-type H3K27ac ChIP-seq data and peaks with ROSE (50).

RNA-seq data were mapped to mouse reference genome mm10 with STAR (v2.7.3a) (51) and quantified with Cufflinks (v2.2.1) (52).

Hi-TrAC raw data were pre-processed into highly-quality non-redundant paired-end tags (PETs) for reference genome mm10 with tracPre2.py in the cLoops2 package. The cLoops2 estSim module performed correlation analysis among replicates, and the cLoops2 estRes module carried out resolution estimations of pooled PETs. The cLoops2 callDomains module called active sub-TADs with parameters of -bs 1000 -ws 25000 -mcut 1000000 -strict. Significantly decreased domains comparing wild-type and *Mll4* knockout cells were called in the same way as human data with the same segregation score cutoff of 1.

### Data visualizations

Most of the tracks were shown by the cLoops2 plot module. Other plots were generated by the matplotlib (53) and seaborn (54) with in-house code.

### Code available

Segregation score based domain-calling algorithm was coded as the cLoops2 callDomains module, domain aggregation analysis was coded in the cLoops2 agg module, domain quantification was coded in the cLoops2 quant module, and insulation score calculation was summarized as getIS.py in the cLoops2 package as a script, all these codes are available at : https://github.com/YaqiangCao/cLoops2.

## DATA AVAILABILITY

Sequencing data generated in this study have been deposited to the Gene Expression Omnibus database with the accession of GSE20808. Visualization tracks are available through the WashU Epigenome Browser with session bundle id: 1321f050-fec0-11ec-882d-4588547110d7.

## ACKNOWLEDGEMENTS

This work utilized the computational resources of the NIH HPC Biowulf cluster (http://hpc.nih.gov). The work was supported by the Division of Intramural Research, National Heart, Lung, and Blood Institute and the 4DN Transformative Collaborative Project Award (A-0066) (KZ).

## AUTHOR CONTRIBUTIONS

Y.C and K.Z. conceived the project. Y.C. analyzed the data. S.L., K.C. contributed to the experiments and Q.T. contributed to the experimental design. Y.C., S.L., and K.Z. wrote the paper. All authors contributed to data interpretation.

## References

1. Marchal, C., Sima, J. and Gilbert, D.M. (2019) Control of DNA replication timing in the 3D genome. Nat Rev Mol Cell Biol, 20, 721–737.

2. Pope, B.D., Ryba, T., Dileep, V., Yue, F., Wu, W., Denas, O., Vera, D.L., Wang, Y., Hansen, R.S., Canfield, T.K. et al. (2014) Topologically associating domains are stable units of replication-timing regulation. Nature, 515, 402–405.

3. Zhang, Y., Zhang, X., Ba, Z., Liang, Z., Dring, E.W., Hu, H., Lou, J., Kyritsis, N., Zurita, J., Shamim, M.S. et al. (2019) The fundamental role of chromatin loop extrusion in physiological V(D)J recombination. Nature, 573, 600–604.

4. Qiu, X., Ma, F., Zhao, M., Cao, Y., Shipp, L., Liu, A., Dutta, A., Singh, A., Braikia, F.Z., De, S. et al. (2020) Altered 3D chromatin structure permits inversional recombination at the IgH locus. Sci Adv, 6, eaaz8850.

5. Hill, L., Ebert, A., Jaritz, M., Wutz, G., Nagasaka, K., Tagoh, H., Kostanova-Poliakova, D., Schindler, K., Sun, Q., Bonelt, P. et al. (2020) Wapl repression by Pax5 promotes V gene recombination by Igh loop extrusion. Nature.

6. Zheng, H. and Xie, W. (2019) The role of 3D genome organization in development and cell differentiation. Nat Rev Mol Cell Biol, 20, 535–550.

7. Stadhouders, R., Filion, G.J. and Graf, T. (2019) Transcription factors and 3D genome conformation in cell-fate decisions. Nature, 569, 345–354.

8. Du, Z., Zheng, H., Huang, B., Ma, R., Wu, J., Zhang, X., He, J., Xiang, Y., Wang, Q., Li, Y. et al. (2017) Allelic reprogramming of 3D chromatin architecture during early mammalian development. Nature, 547, 232–235.

9. Ke, Y., Xu, Y., Chen, X., Feng, S., Liu, Z., Sun, Y., Yao, X., Li, F., Zhu, W., Gao, L. et al. (2017) 3D Chromatin Structures of Mature Gametes and Structural Reprogramming during Mammalian Embryogenesis. Cell, 170, 367–381 e320.

10. Wang, H., Han, M. and Qi, L.S. (2021) Engineering 3D genome organization. Nat Rev Genet, 22, 343–360.

11. Lupianez, D.G., Kraft, K., Heinrich, V., Krawitz, P., Brancati, F., Klopocki, E., Horn, D., Kayserili, H., Opitz, J.M., Laxova, R. et al. (2015) Disruptions of topological chromatin domains cause pathogenic rewiring of gene-enhancer interactions. Cell, 161, 1012–1025.

12. Hnisz, D., Weintraub, A.S., Day, D.S., Valton, A.L., Bak, R.O., Li, C.H., Goldmann, J., Lajoie, B.R., Fan, Z.P., Sigova, A.A. et al. (2016) Activation of proto-oncogenes by disruption of chromosome neighborhoods. Science, 351, 1454–1458.

13. Flavahan, W.A., Drier, Y., Liau, B.B., Gillespie, S.M., Venteicher, A.S., Stemmer-Rachamimov, A.O., Suva, M.L. and Bernstein, B.E. (2016) Insulator dysfunction and oncogene activation in IDH mutant gliomas. Nature, 529, 110–114.

14. Jerkovic, I. and Cavalli, G. (2021) Understanding 3D genome organization by multidisciplinary methods. Nat Rev Mol Cell Biol, 22, 511–528.

15. Kempfer, R. and Pombo, A. (2020) Methods for mapping 3D chromosome architecture. Nat Rev Genet, 21, 207–226.

16. Lakadamyali, M. and Cosma, M.P. (2020) Visualizing the genome in high resolution challenges our textbook understanding. Nat Methods, 17, 371–379.

17. Lieberman-Aiden, E., van Berkum, N.L., Williams, L., Imakaev, M., Ragoczy, T., Telling, A., Amit, I., Lajoie, B.R., Sabo, P.J., Dorschner, M.O. et al. (2009) Comprehensive mapping of long-range interactions reveals folding principles of the human genome. Science, 326, 289–293.

18. Dixon, J.R., Selvaraj, S., Yue, F., Kim, A., Li, Y., Shen, Y., Hu, M., Liu, J.S. and Ren, B. (2012) Topological domains in mammalian genomes identified by analysis of chromatin interactions. Nature, 485, 376–380.

19. Sexton, T., Yaffe, E., Kenigsberg, E., Bantignies, F., Leblanc, B., Hoichman, M., Parrinello, H., Tanay, A. and Cavalli, G. (2012) Three-dimensional folding and functional organization principles of the Drosophila genome. Cell, 148, 458–472.

20. Nora, E.P., Lajoie, B.R., Schulz, E.G., Giorgetti, L., Okamoto, I., Servant, N., Piolot, T., van Berkum, N.L., Meisig, J., Sedat, J. et al. (2012) Spatial partitioning of the regulatory landscape of the X-inactivation centre. Nature, 485, 381–385.

21. Rao, Suhas S.P., Huntley, Miriam H., Durand, Neva C., Stamenova, Elena K., Bochkov, Ivan D., Robinson, James T., Sanborn, Adrian L., Machol, I., Omer, Arina D., Lander, Eric S. et al. (2014) A 3D Map of the Human Genome at Kilobase Resolution Reveals Principles of Chromatin Looping. Cell, 159, 1665–1680.

22. Hsieh, T.S., Cattoglio, C., Slobodyanyuk, E., Hansen, A.S., Rando, O.J., Tjian, R. and Darzacq, X. (2020) Resolving the 3D Landscape of Transcription-Linked Mammalian Chromatin Folding. Mol Cell.

23. Krietenstein, N., Abraham, S., Venev, S.V., Abdennur, N., Gibcus, J., Hsieh, T.S., Parsi, K.M., Yang, L., Maehr, R., Mirny, L.A. et al. (2020) Ultrastructural Details of Mammalian Chromosome Architecture. Mol Cell.

24. Tang, Z., Luo, Oscar J., Li, X., Zheng, M., Zhu, Jacqueline J., Szalaj, P., Trzaskoma, P., Magalska, A., Wlodarczyk, J., Ruszczycki, B. et al. (2015) CTCF-Mediated Human 3D Genome Architecture Reveals Chromatin Topology for Transcription. Cell, 163, 1611–1627.

25. Beagrie, R.A., Scialdone, A., Schueler, M., Kraemer, D.C.A., Chotalia, M., Xie, S.Q., Barbieri, M., de Santiago, I., Lavitas, L.-M., Branco, M.R. et al. (2017) Complex multi-enhancer contacts captured by genome architecture mapping. Nature, 543, 519–524.

26. Quinodoz, S.A., Ollikainen, N., Tabak, B., Palla, A., Schmidt, J.M., Detmar, E., Lai, M.M., Shishkin, A.A., Bhat, P., Takei, Y. et al. (2018) Higher-Order Inter-chromosomal Hubs Shape 3D Genome Organization in the Nucleus. Cell, 174, 744–757 e724.

27. Lai, B., Tang, Q., Jin, W., Hu, G., Wangsa, D., Cui, K., Stanton, B.Z., Ren, G., Ding, Y., Zhao, M. et al. (2018) Trac-looping measures genome structure and chromatin accessibility. Nat Methods, 15, 741–747.

28. Crane, E., Bian, Q., McCord, R.P., Lajoie, B.R., Wheeler, B.S., Ralston, E.J., Uzawa, S., Dekker, J. and Meyer, B.J. (2015) Condensin-driven remodelling of X chromosome topology during dosage compensation. Nature, 523, 240–244.

29. Liu, C., Cheng, Y.J., Wang, J.W. and Weigel, D. (2017) Prominent topologically associated domains differentiate global chromatin packing in rice from Arabidopsis. Nat Plants, 3, 742–748.

30. Phillips-Cremins, J.E., Sauria, M.E., Sanyal, A., Gerasimova, T.I., Lajoie, B.R., Bell, J.S., Ong, C.T., Hookway, T.A., Guo, C., Sun, Y. et al. (2013) Architectural protein subclasses shape 3D organization of genomes during lineage commitment. Cell, 153, 1281–1295.

31. Schwarzer, W., Abdennur, N., Goloborodko, A., Pekowska, A., Fudenberg, G., Loe-Mie, Y., Fonseca, N.A., Huber, W., Haering, C.H., Mirny, L. et al. (2017) Two independent modes of chromatin organization revealed by cohesin removal. Nature, 551, 51–56.

32. Haarhuis, J.H.I., van der Weide, R.H., Blomen, V.A., Yanez-Cuna, J.O., Amendola, M., van Ruiten, M.S., Krijger, P.H.L., Teunissen, H., Medema, R.H., van Steensel, B. et al. (2017) The Cohesin Release Factor WAPL Restricts Chromatin Loop Extension. Cell, 169, 693–707 e614.

33. Davidson, I.F. and Peters, J.M. (2021) Genome folding through loop extrusion by SMC complexes. Nat Rev Mol Cell Bio, 22, 445–464.

34. Szabo, Q., Bantignies, F. and Cavalli, G. (2019) Principles of genome folding into topologically associating domains. Science Advances, 5.

35. Beagan, J.A. and Phillips-Cremins, J.E. (2020) On the existence and functionality of topologically associating domains. Nat Genet, 52, 8–16.

36. Ou, H.D., Phan, S., Deerinck, T.J., Thor, A., Ellisman, M.H. and O’Shea, C.C. (2017) ChromEMT: Visualizing 3D chromatin structure and compaction in interphase and mitotic cells. Science, 357.

37. Liu, S., Cao, Y., Cui, K., Tang, Q. and Zhao, K. (2022) Hi-TrAC reveals fine-scale chromatin structures organized by transcription factors. bioRxiv, 2022.2006.2001.494329.

38. Cao, Y., Liu, S., Ren, G., Tang, Q. and Zhao, K. (2022) cLoops2: a full-stack comprehensive analytical tool for chromatin interactions. Nucleic Acids Res, 50, 57–71.

39. Frankish, A., Diekhans, M., Ferreira, A.M., Johnson, R., Jungreis, I., Loveland, J., Mudge, J.M., Sisu, C., Wright, J., Armstrong, J. et al. (2019) GENCODE reference annotation for the human and mouse genomes. Nucleic Acids Res, 47, D766–D773.

40. Khan, A. and Zhang, X. (2016) dbSUPER: a database of super-enhancers in mouse and human genome. Nucleic Acids Res, 44, D164–171.

41. Abadi, M., Agarwal, A., Barham, P., Brevdo, E., Chen, Z., Citro, C., Corrado, G.S., Davis, A., Dean, J. and Devin, M. (2016) Tensorflow: Large-scale machine learning on heterogeneous distributed systems. arXiv preprint 1603.04467.

42. Pedregosa, F., Varoquaux, G., Gramfort, A., Michel, V., Thirion, B., Grisel, O., Blondel, M., Prettenhofer, P., Weiss, R. and Dubourg, V. (2011) Scikit-learn: Machine learning in Python. the Journal of machine Learning research, 12, 2825–2830.

43. Heinz, S., Benner, C., Spann, N., Bertolino, E., Lin, Y.C., Laslo, P., Cheng, J.X., Murre, C., Singh, H. and Glass, C.K. (2010) Simple Combinations of Lineage-Determining Transcription Factors Prime cis-Regulatory Elements Required for Macrophage and B Cell Identities. Molecular Cell, 38, 576–589.

44. Placek, K., Hu, G., Cui, K., Zhang, D., Ding, Y., Lee, J.E., Jang, Y., Wang, C., Konkel, J.E., Song, J. et al. (2017) MLL4 prepares the enhancer landscape for Foxp3 induction via chromatin looping. Nat Immunol, 18, 1035–1045.

45. Barski, A., Cuddapah, S., Cui, K., Roh, T.Y., Schones, D.E., Wang, Z., Wei, G., Chepelev, I. and Zhao, K. (2007) High-resolution profiling of histone methylations in the human genome. Cell, 129, 823–837.

46. Wei, G., Abraham, B.J., Yagi, R., Jothi, R., Cui, K., Sharma, S., Narlikar, L., Northrup, D.L., Tang, Q., Paul, W.E. et al. (2011) Genome-wide analyses of transcription factor GATA3-mediated gene regulation in distinct T cell types. Immunity, 35, 299–311.

47. Picelli, S., Faridani, O.R., Bjorklund, A.K., Winberg, G., Sagasser, S. and Sandberg, R. (2014) Full-length RNA-seq from single cells using Smart-seq2. Nat Protoc, 9, 171–181.

48. Langmead, B. and Salzberg, S.L. (2012) Fast gapped-read alignment with Bowtie 2. Nature Methods, 9, 357–U354.

49. Ramirez, F., Ryan, D.P., Gruning, B., Bhardwaj, V., Kilpert, F., Richter, A.S., Heyne, S., Dundar, F. and Manke, T. (2016) deepTools2: a next generation web server for deep-sequencing data analysis. Nucleic Acids Res, 44, W160–165.

50. Whyte, W.A., Orlando, D.A., Hnisz, D., Abraham, B.J., Lin, C.Y., Kagey, M.H., Rahl, P.B., Lee, T.I. and Young, R.A. (2013) Master Transcription Factors and Mediator Establish Super-Enhancers at Key Cell Identity Genes. Cell, 153, 307–319.

51. Dobin, A., Davis, C.A., Schlesinger, F., Drenkow, J., Zaleski, C., Jha, S., Batut, P., Chaisson, M. and Gingeras, T.R. (2013) STAR: ultrafast universal RNA-seq aligner. Bioinformatics, 29, 15–21.

52. Trapnell, C., Hendrickson, D.G., Sauvageau, M., Goff, L., Rinn, J.L. and Pachter, L. (2013) Differential analysis of gene regulation at transcript resolution with RNA-seq. Nature Biotechnology, 31, 46–+.

53. Hunter, J.D. (2007) Matplotlib: A 2D graphics environment. Computing in science & engineering, 9, 90–95.

54. Waskom, M.L. (2021) Seaborn: statistical data visualization. Journal of Open Source Software, 6, 3021.

55. Olivares-Chauvet, P., Mukamel, Z., Lifshitz, A., Schwartzman, O., Elkayam, N.O., Lubling, Y., Deikus, G., Sebra, R.P. and Tanay, A. (2016) Capturing pairwise and multi-way chromosomal conformations using chromosomal walks. Nature, 540, 296–+.

56. Rouault, J.P., Falette, N., Guehenneux, F., Guillot, C., Rimokh, R., Wang, Q., Berthet, C., Moyret-Lalle, C., Savatier, P., Pain, B. et al. (1996) Identification of BTG2, an antiproliferative p53-dependent component of the DNA damage cellular response pathway. Nat Genet, 14, 482–486.

57. Wang, J.H., Avitahl, N., Cariappa, A., Friedrich, C., Ikeda, T., Renold, A., Andrikopoulos, K., Liang, L.B., Pillai, S., Morgan, B.A. et al. (1998) Aiolos regulates B cell activation and maturation to effector state. Immunity, 9, 543–553.

58. Gong, Y.X., Lazaris, C., Sakellaropoulos, T., Lozano, A., Kambadur, P., Ntziachristos, P., Aifantis, I. and Tsirigos, A. (2018) Stratification of TAD boundaries reveals preferential insulation of super-enhancers by strong boundaries. Nature Communications, 9.

59. Parelho, V., Hadjur, S., Spivakov, M., Leleu, M., Sauer, S., Gregson, H.C., Jarmuz, A., Canzonetta, C., Webster, Z., Nesterova, T. et al. (2008) Cohesins functionally associate with CTCF on mammalian chromosome arms. Cell, 132, 422–433.

60. Wendt, K.S., Yoshida, K., Itoh, T., Bando, M., Koch, B., Schirghuber, E., Tsutsumi, S., Nagae, G., Ishihara, K., Mishiro, T. et al. (2008) Cohesin mediates transcriptional insulation by CCCTC-binding factor. Nature, 451, 796–801.

61. Harrold, C.L., Gosden, M.E., Hanssen, L.L.P., Stolper, R.J., Downes, D.J., Telenius, J.M., Biggs, D., Preece, C., Alghadban, S., Sharpe, J.A. et al. (2020) A functional overlap between actively transcribed genes and chromatin boundary elements. bioRxiv, 2020.2007.2001.182089.

62. Oudelaar, A.M. and Higgs, D.R. (2021) The relationship between genome structure and function. Nat Rev Genet, 22, 154–168.

63. Hnisz, D., Day, D.S. and Young, R.A. (2016) Insulated Neighborhoods: Structural and Functional Units of Mammalian Gene Control. Cell, 167, 1188–1200.

64. Naughton, C., Avlonitis, N., Corless, S., Prendergast, J.G., Mati, I.K., Eijk, P.P., Cockroft, S.L., Bradley, M., Ylstra, B. and Gilbert, N. (2013) Transcription forms and remodels supercoiling domains unfolding large-scale chromatin structures. Nat Struct Mol Biol, 20, 387–395.

65. Szabo, Q., Donjon, A., Jerkovic, I., Papadopoulos, G.L., Cheutin, T., Bonev, B., Nora, E.P., Bruneau, B.G., Bantignies, F. and Cavalli, G. (2020) Regulation of single-cell genome organization into TADs and chromatin nanodomains. Nat Genet, 52, 1151–+.

66. Grubert, F., Srivas, R., Spacek, D.V., Kasowski, M., Ruiz-Velasco, M., Sinnott-Armstrong, N., Greenside, P., Narasimha, A., Liu, Q., Geller, B. et al. (2020) Landscape of cohesin-mediated chromatin loops in the human genome. Nature, 583, 737–743.

67. Mumbach, M.R., Satpathy, A.T., Boyle, E.A., Dai, C., Gowen, B.G., Cho, S.W., Nguyen, M.L., Rubin, A.J., Granja, J.M., Kazane, K.R. et al. (2017) Enhancer connectome in primary human cells identifies target genes of disease-associated DNA elements. Nat Genet, 49, 1602–1612.

68. Alipour, E. and Marko, J.F. (2012) Self-organization of domain structures by DNA-loop-extruding enzymes. Nucleic Acids Res, 40, 11202–11212.

69. Dowen, J.M., Fan, Z.P., Hnisz, D., Ren, G., Abraham, B.J., Zhang, L.N., Weintraub, A.S., Schujiers, J., Lee, T.I., Zhao, K. et al. (2014) Control of cell identity genes occurs in insulated neighborhoods in mammalian chromosomes. Cell, 159, 374–387.

70. Consortium, E.P. (2012) An integrated encyclopedia of DNA elements in the human genome. Nature, 489, 57–74.

71. Yan, J., Chen, S.A., Local, A., Liu, T., Qiu, Y., Dorighi, K.M., Preissl, S., Rivera, C.M., Wang, C., Ye, Z. et al. (2018) Histone H3 lysine 4 monomethylation modulates long-range chromatin interactions at enhancers. Cell Res, 28, 204–220.

72. Karmodiya, K., Krebs, A.R., Oulad-Abdelghani, M., Kimura, H. and Tora, L. (2012) H3K9 and H3K14 acetylation co-occur at many gene regulatory elements, while H3K14ac marks a subset of inactive inducible promoters in mouse embryonic stem cells. Bmc Genomics, 13.

73. Kim, Y.J., Sekiya, F., Poulin, B., Bae, Y.S. and Rhee, S.G. (2004) Mechanism of B-cell receptor-induced phosphorylation and activation of phospholipase C-gamma2. Mol Cell Biol, 24, 9986–9999.

74. Lilly, M., Le, T., Holland, P. and Hendrickson, S.L. (1992) Sustained Expression of the Pim-1 Kinase Is Specifically Induced in Myeloid Cells by Cytokines Whose Receptors Are Structurally Related. Oncogene, 7, 727–732.

75. Rao, S.S.P., Huang, S.C., Glenn St Hilaire, B., Engreitz, J.M., Perez, E.M., Kieffer-Kwon, K.R., Sanborn, A.L., Johnstone, S.E., Bascom, G.D., Bochkov, I.D. et al. (2017) Cohesin Loss Eliminates All Loop Domains. Cell, 171, 305–320 e324.

76. McCormack, M.P., Young, L.F., Vasudevan, S., de Graaf, C.A., Codrington, R., Rabbitts, T.H., Jane, S.M. and Curtis, D.J. (2010) The Lmo2 Oncogene Initiates Leukemia in Mice by Inducing Thymocyte Self-Renewal. Science, 327, 879–883.

77. Lee, J.E., Wang, C.C., Xu, S.L.Y., Cho, Y.W., Wang, L.F., Feng, X.S., Baldridge, A., Sartorelli, V., Zhuang, L.N., Peng, W.Q. et al. (2013) H3K4 mono- and di-methyltransferase MLL4 is required for enhancer activation during cell differentiation. Elife, 2.

78. Dong, C. (2021) Cytokine Regulation and Function in T Cells. Annu Rev Immunol, 39, 51–76.

79. Alunno, A., Bartoloni, E., Bistoni, O., Nocentini, G., Ronchetti, S., Caterbi, S., Valentini, V., Riccardi, C. and Gerli, R. (2012) Balance between Regulatory T and Th17 Cells in Systemic Lupus Erythematosus: The Old and the New. Clin Dev Immunol.

80. Shah, K., Lee, W.W., Lee, S.H., Kim, S.H., Kang, S.W., Craft, J. and Kang, I. (2010) Dysregulated balance of Th17 and Th1 cells in systemic lupus erythematosus (vol 12, pg 402, 2010). Arthritis Res Ther, 12.

81. Dolff, S., Bijl, M., Huitema, M.G., Limburg, P.C., Kallenberg, C.G. and Abdulahad, W.H. (2011) Disturbed Th1, Th2, Th17 and T(reg) balance in patients with systemic lupus erythematosus. Clin Immunol, 141, 197–204.

82. Chen, X., Zhang, M., Liao, M., Graner, M.W., Wu, C., Yang, Q., Liu, H. and Zhou, B. (2010) Reduced Th17 response in patients with tuberculosis correlates with IL-6R expression on CD4+ T Cells. Am J Respir Crit Care Med, 181, 734–742.

83. Jones, G.W., McLoughlin, R.M., Hammond, V.J., Parker, C.R., Williams, J.D., Malhotra, R., Scheller, J., Williams, A.S., Rose-John, S., Topley, N. et al. (2010) Loss of CD4+ T cell IL-6R expression during inflammation underlines a role for IL-6 trans signaling in the local maintenance of Th17 cells. J Immunol, 184, 2130–2139.

84. Flyamer, I.M., Gassler, J., Imakaev, M., Brandao, H.B., Ulianov, S.V., Abdennur, N., Razin, S.V., Mirny, L.A. and Tachibana-Konwalski, K. (2017) Single-nucleus Hi-C reveals unique chromatin reorganization at oocyte-to-zygote transition. Nature, 544, 110–+.

85. Nagano, T., Lubling, Y., Varnai, C., Dudley, C., Leung, W., Baran, Y., Mendelson Cohen, N., Wingett, S., Fraser, P. and Tanay, A. (2017) Cell-cycle dynamics of chromosomal organization at single-cell resolution. Nature, 547, 61–67.

86. Stevens, T.J., Lando, D., Basu, S., Atkinson, L.P., Cao, Y., Lee, S.F., Leeb, M., Wohlfahrt, K.J., Boucher, W., O’Shaughnessy-Kirwan, A. et al. (2017) 3D structures of individual mammalian genomes studied by single-cell Hi-C. Nature, 544, 59–+.

87. Consortium, E.P., Moore, J.E., Purcaro, M.J., Pratt, H.E., Epstein, C.B., Shoresh, N., Adrian, J., Kawli, T., Davis, C.A., Dobin, A. et al. (2020) Expanded encyclopaedias of DNA elements in the human and mouse genomes. Nature, 583, 699–710.

88. Djebali, S., Davis, C.A., Merkel, A., Dobin, A., Lassmann, T., Mortazavi, A., Tanzer, A., Lagarde, J., Lin, W., Schlesinger, F. et al. (2012) Landscape of transcription in human cells. Nature, 489, 101–108.

