## Supplemental Information for "Hi-TrAC reveals fractal nesting of super-enhancers"

### Figure S1

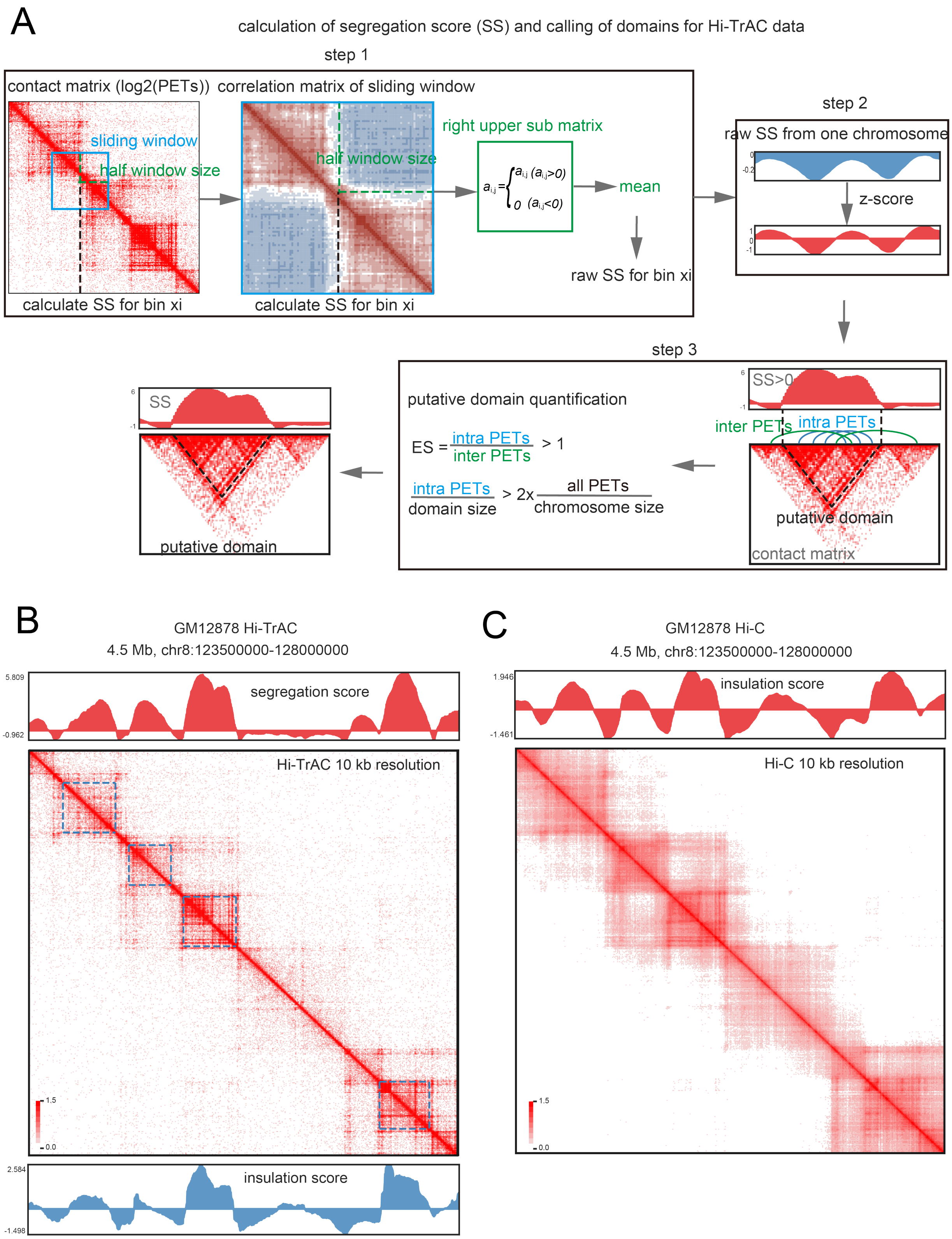
A
calculation of segregation score (SS) and calling of domains for Hi-TrAC data
step 1
contact matrix (log2(PETs))
correlation matrix of sliding window
sliding window
half window size
right upper sub matrix
half window size
mean
raw SS for bin xi
calculate SS for bin xi
calculate SS for bin xi
step 2
raw SS from one chromosome
z-score
step 3
putative domain quantification
SS>0
inter PETs
intra PETs
ES = intra PETs / inter PETs > 1
intra PETs / domain size > 2x all PETs / chromosome size
putative domain
contact matrix
SS
putative domain
B
GM12878 Hi-TrAC
4.5 Mb, chr8:123500000-128000000
5.809
-0.962
segregation score
Hi-TrAC 10 kb resolution
1.5
0.0
2.584
-1.498
insulation score
C
GM12878 Hi-C
4.5 Mb, chr8:123500000-128000000
1.946
-1.461
insulation score
Hi-C 10 kb resolution
1.5
0.0

**Figure S1. Segregation score calculation and domain calling from Hi-TrAC data.**

(A) Computational scheme of segregation score calculation and putative domain calling from Hi-TrAC data. Details are described in the **Methods**.

(B) Examples of domains called from Hi-TrAC data of GM12878 cells at a 10 kb resolution. Domains were marked as blue dotted frames on the heatmap, and segregation scores were shown as the top red track. Insulation scores calculated with the same parameters (resolution and sliding window size) as the segregation scores from Hi-TrAC data were shown as the blue bottom track.

(C) Example of interaction heatmap of GM12878 in situ Hi-C data from Rao et al. 2014. The genomic region is the same as panel (B). The Hi-C insulation scores were calculated by getIS.py in the cLoops2 package with the parameters of -bs 10000 -s 500000.

Figure S2

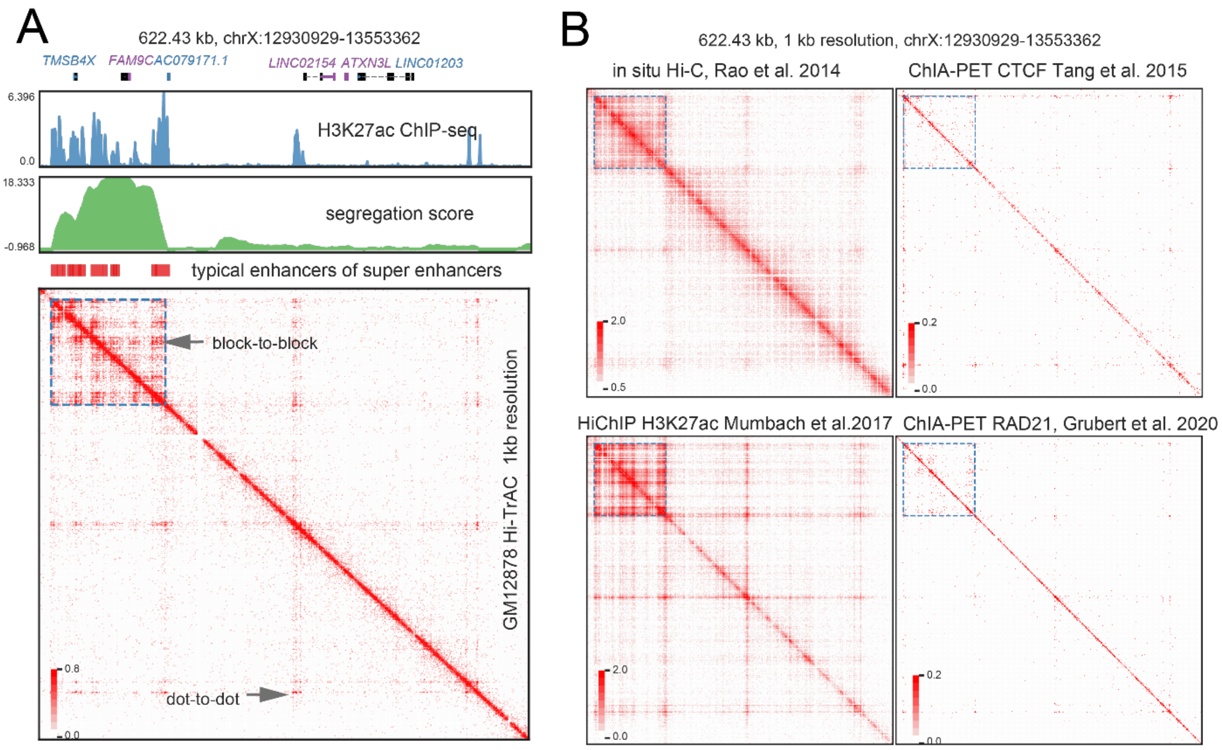


**Figure S2. Comparison of fine-scale structures of super-enhancers detected by various techniques.**

(A) An example of interactions detected by Hi-TrAC at a super-enhancer containing sub-TAD and nearby regions in GM12878.

(B) Interaction matrix heatmaps from the data generated by other chromatin architecture mapping technologies for the same region as in Panel (A).

### Figure S3

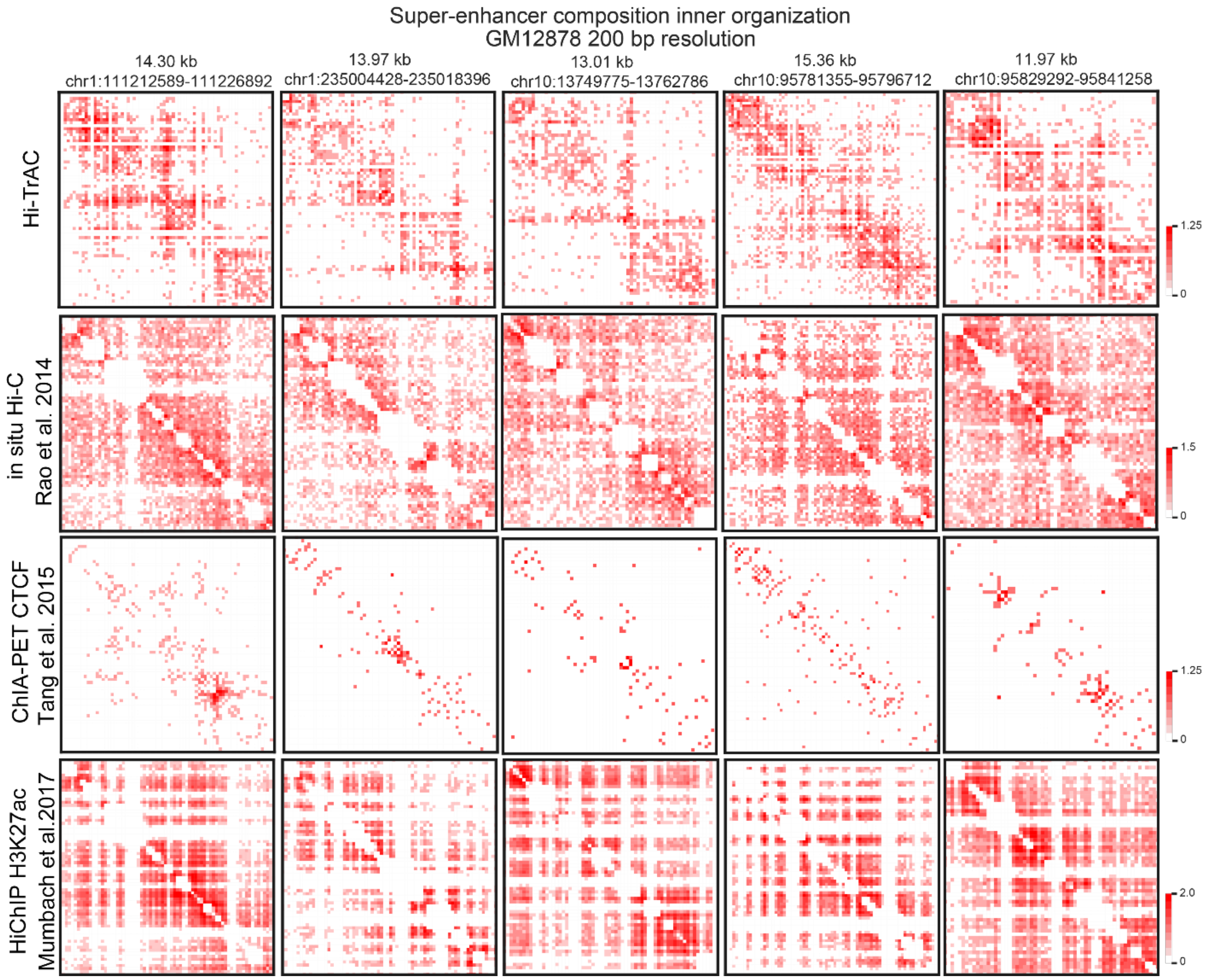


**Figure S3. Comparison of super-enhancers' inner structures detected by various techniques.** The interaction matrix heatmaps at 200 bp resolution for randomly selected five super-enhancers are shown to compare the data from Hi-TrAC with other representative technologies. Only Hi-TrAC detects the explicit inner structures.

### Figure S4

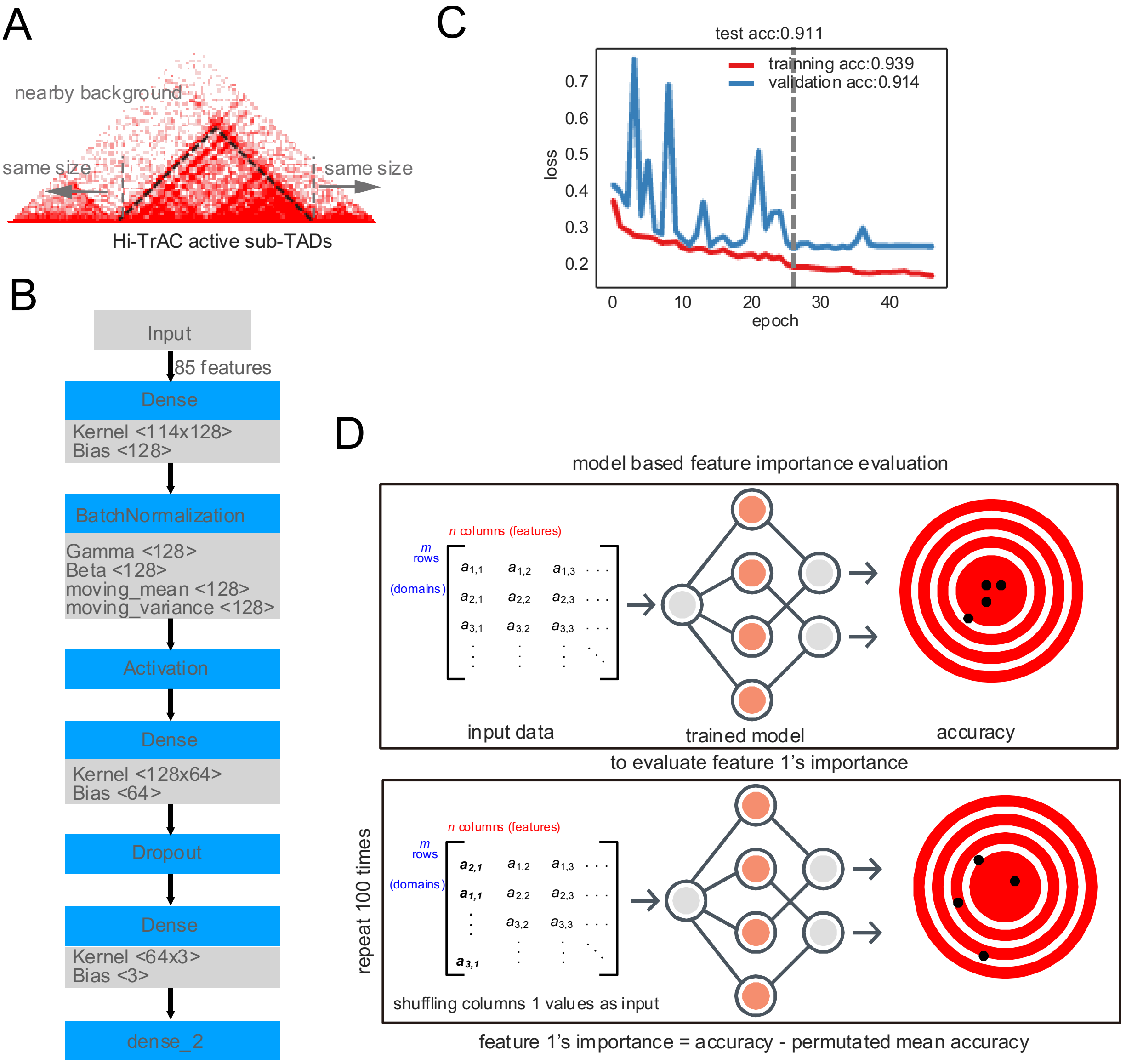


**Figure S4. Classification model for ChIP-seq data to distinguish Hi-TrAC active sub-TADs against background regions.**

(A) Definition of background regions for comparison with Hi-TrAC active sub-TADs. Background regions were defined as the same-sized flanking regions to sub-TADs but not overlapped with any of them.

(B) Scheme plot of a simple deep learning model for classifying Hi-TrAC sub-TADs and background regions based on 85 shared chromatin features between GM12878 and K562 (binding of transcription factors and histone modifications) quantified from ENCODE ChIP-seq data.

(C) Training history of the simple deep learning model. The training model at epoch 27 was saved for usage. The consistency losses of training and validation data indicate that the model is not overfitted. The similar accuracy of the final test dataset to that of training and validation data indicates that the model captures the authentic latent features for classification.

(D) Scheme plot of model-based feature importance evaluation. The estimation for feature importance was performed for each factor by shuffling the RPKM values for all regions 100 times. Then the mean value of decreased accuracy was used as the feature importance.

### Figure S5

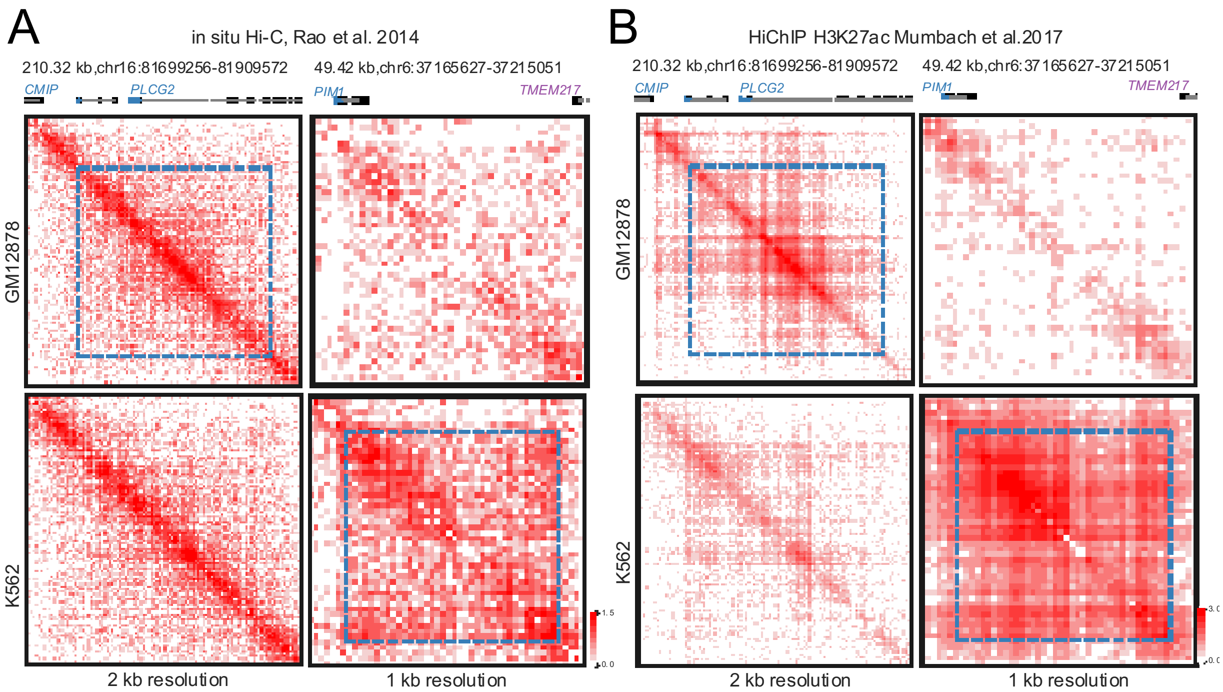


**Figure S5. Examples of validation of Hi-TrAC cell-specific active sub-TADs by Hi-C and H3K27ac HiChIP.**

(A) Examples of validation of a GM12878 and K562 specific sub-TAD by Hi-C data.

(B) Examples of validation of a GM12878 and K562 specific sub-TAD by H3K27ac HiChIP data.

### Figure S6

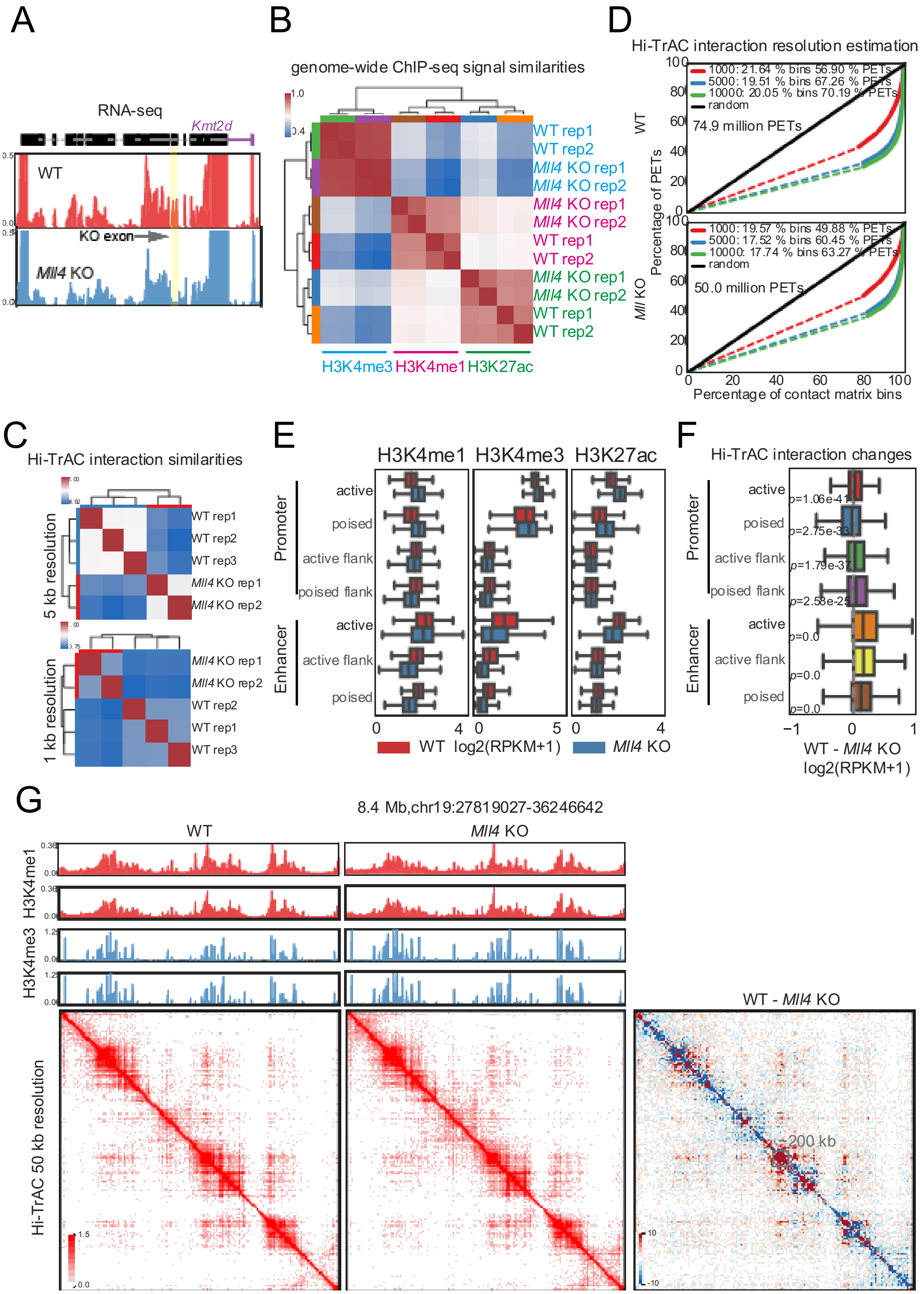
A
RNA-seq
Kmt2d
WT
KO exon
Mll4 KO
B
genome-wide ChIP-seq signal similarities
WT rep1
WT rep2
Mll4 KO rep1
Mll4 KO rep2
H3K4me3
H3K4me1
H3K27ac
D
Hi-TrAC interaction resolution estimation
1000: 21.64 % bins 56.90 % PETs
5000: 19.51 % bins 67.26 % PETs
10000: 20.05 % bins 70.19 % PETs
random
74.9 million PETs
1000: 19.57 % bins 49.88 % PETs
5000: 17.52 % bins 60.45 % PETs
10000: 17.74 % bins 63.27 % PETs
50.0 million PETs
Percentage of PETs
Percentage of contact matrix bins
C
Hi-TrAC interaction similarities
5 kb resolution
1 kb resolution
WT rep3
E
Promoter
Enhancer
active
poised
active flank
poised flank
log2(RPKM+1)
F
Hi-TrAC interaction changes
p=1.06e-41
p=2.75e-33
p=1.79e-37
p=2.58e-25
p=0.0
WT - Mll4 KO
log2(RPKM+1)
G
8.4 Mb,chr19:27819027-36246642
Hi-TrAC 50 kb resolution
200 kb

**Figure S6. H3K4me1 and interaction at putative enhancers are decreased after deletion of *Mll4* in mouse Th17 cells.**

(A) Validation of *Mll4*'s exon conditional knockout by RNA-seq data.

(B) Sample-wise genome-wide similarity analysis for ChIP-seq data.

(C) Sample-wise genome-wide similarity analysis for Hi-TrAC interactions at 5 kb and 1 kb resolution, respectively.

(D) Resolution estimation analysis for pooled Hi-TrAC data.

(E) Distribution of ChIP-seq signals at genomic segments. Segments were defined from overlaps of H3K4me1, H3K4me3, and H3K27ac peaks. Briefly, H3K4m3 overlapped H3K27ac peaks at the transcription start site (TSS) were assigned as active TSS first. H3K4me3 peaks at TSS with H3K27ac peaks were assigned as poised TSS. Then H3K27ac peaks at TSS distal regions were assigned as active enhancers. H3K4me1 only peaks at TSS distal region were assigned as poised enhancers. Finally, other parts of H3K4me1 peaks overlapped with above segments but not totally covered as the flank regions.

(F) Distribution of changes of Hi-TrAC interaction densities at genomic segments. The Wilcoxon signed-rank test *P*-values were annotated in the boxplot plot.

(G) Example of interaction changes comparing wild-type and *Mll4*-deleted Th17 cells. Gray square annotates a region about 200 kb size showing decreased interaction densities.
